# Purkinje cells in Crus I and II encode the visual stimulus and the impending choice as monkeys learn a reinforcement based visuomotor association task

**DOI:** 10.1101/2024.09.13.612926

**Authors:** Anna E. Ipata, Valeria Fascianelli, Chris I. De Zeeuw, Naveen Sendhilnathan, Stefano Fusi, Michael E. Goldberg

**Affiliations:** Dept. of Neuroscience, Columbia University, New York, NY, USA; Mahoney Center for Brain and Behavior Research, Columbia University, New York, NY, USA; Center for Theoretical Neuroscience, Columbia University, NY, USA; Dept. of Neuroscience, Erasmus MC, Rotterdam, The Netherlands; Netherlands Institute for Neuroscience, Royal Dutch Academy of Arts and Science (KNAW), Amsterdam, The Netherlands; Kavli Institute for Brain Science, Columbia University, New York, NY, USA; Zuckerman Mind, Brain, and Behavior Institute, Columbia University, New York, NY, USA; Dept. of Neurology, Psychiatry, and Ophthalmology, Columbia University College of Physicians and Surgeons, New York, NY, USA

## Abstract

Visuomotor association involves linking an arbitrary visual cue to a well-learned movement. Transient inactivation of Crus I/II impairs primates’ ability to learn new associations and delays motor responses without affecting the kinematics of the movement. The simple spikes of Purkinje cells in the Crus regions signal cognitive errors as monkeys learn to associate specific fractal stimuli with movements of the left or right hand. Here we show that as learning progresses, the simple spike activity of individual neurons becomes more selective for stimulus-response associations, with selectivity developing closer to the appearance of visual stimuli. Initially, most neurons respond to both associations, irrespective of the identity of the stimulus and the associated movement, but as learning advances, more neurons distinguish between specific stimulus-hand associations. Using a linear decoder, it was found that in early learning stages, the visual stimulus can be decoded only when the choice can also be decoded. As learning improves, the visual stimulus is decoded earlier than the choice. A simple model can replicate the observed simple spike signals and the monkeys’ behavior in both the early and late learning stages.

## Introduction

Although the cerebellum has traditionally been associated primarily with motor control and learning^1^, recent studies show that the posterolateral cerebellum is critical for visuomotor association learning. When monkeys perform an overtrained visuomotor association task (OT) in which they associate one of two familiar stimuli with a left hand movement and the other with a right hand movement, the great majority of simple spikes (SSs) of Purkinje cells (PCells) in Crus I and Crus II discharge in the interval between the stimulus appearance and the response movement, but are not sensitive to the presence or absence of reward in the prior trial. When the stimulus switches to a pair of fractal stimuli that the monkey has never seen before, the SSs exhibit, during this period of early learning (EL), an error signal based on the outcome of the prior trial^2^ even though the kinematics of the response movement are unchanged. This error signal decreases as the monkey learns the task. Accordingly, transient inactivation of Crus I and II in the cerebellum significantly impairs the ability of primates to learn the association, while the kinematics of the motor response itself remain unaffected^3^. Instead, when the monkeys respond to overtrained stimuli, inactivation of Crus I and II does not affect the monkeys’ ability to remember the visuomotor associations. Although transient inactivation of Crus I and II never affected the kinematics of the motor responses, i.e., neither in the EL nor in the OT stage, it does prevent the shortening of the response latencies that normally come with training^3^.

In a preliminary analysis of the data, we found that, across the population, there was no average difference in the response of SSs to the monkey’s hand response or stimulus in the peristimulus epoch time^2^. Here, we reanalyzed these data, using trial-by-trial analysis of the neuronal responses. We found that SSs of individual PCells responded at different times after the stimulus appeared, tiling the entire interval between stimulus appearance and response. As learning progressed, the number of PCells that responded before the impending movement increased, and the time between the neural response and the beginning of the movement decreased. Furthermore, the majority of cells responded with significantly different response intensities when the monkey responded with the left as opposed to the right hand. Similarly, more cells showed this difference in response during OT than during EL.

When we applied a linear decoding method to the population activity, we found that SSs had information about both the stimulus presented and the monkey’s hand response. Decoding was more accurate during OT than EL and occurred earlier relative to stimulus appearance and longer before the response. We developed two models to help us better understand these data. First, we hypothesized that the visual inputs to the cerebellum from prefrontal cortex became stronger as the learning progressed. Second, we trained a simple feedforward neural network to perform the learning tasks, using the same assumptions on the dynamics of the input signals to PCells. Both models duplicated the experimental results.

## Results

The current study is a reanalysis of previously published data^2^. Two monkeys were trained for several months to perform a visuomotor association task where they had to associate one visual stimulus with a left-hand movement and another visual stimulus with a right-hand movement. The monkeys began each trial by placing their left and right hand on a bar on the left and right side, respectively (Figure 1A), after which a white square, the fixation point, appeared at the center of the screen. Monkeys had to fixate on the white square for an interval between 100ms and 800ms, and then a green or pink square appeared, instructing them to release the left hand or the right hand, respectively. We trained the monkeys on the overtrained (OT) colored squares until they reached performance accuracy close to 100%. We began each experiment with 20-30 trials using the overtrained stimuli, making sure that the monkeys performed with a success rate of at least 90%. We then switched the stimuli to fractal pictures^4^ that the monkeys had never seen before (New Task, NT). We used logistic regression to estimate the time course of the monkeys’ performance during learning the new associations. (Figure 1B). We divided the trials into two distinct categories: Early Learning (EL), i.e., all trials with a probability of being correct less than 75%, and Late Learning (LL), i.e., all trials with a probability of being correct higher than 75%.

### Purkinje Cells respond at different times during the trial to the Stimulus-Response association

We recorded the activity of SSs in 80 Purkinje cells (PCells) in the right posterolateral cerebellum, predominantly in Crus I and II in two monkeys (Figure 1C). We only analyzed cells that were stable throughout the entire session. We analyzed correct and error trials separately. Because of the structure of OT and NT tasks, the stimulus –stimulus 1 and stimulus 2– and response –right and left– are perfectly correlated, we define the response of the neurons after stimulus onset as the Stimulus-Response left (SR left) activity, when monkey released the left bar, and Stimulus-response right (SR right) activity, when monkeys released the right bar. Figure 2A shows the responses of 3 representative neurons in OT (left column), LL (middle) and EL (right) for correct trials (see Supplementary Figure 1 for more example neurons). In the overtrained task, neuron N1 showed an increase in firing rate immediately before movement, more prominently during SR left than SR right trials; Neuron N2 displayed inhibition following stimulus onset followed by a gradual increase in activity leading up to the movement for both SR left and SR right trials; Neuron N3 demonstrated a decrease in activity after movement onset in both SR left and SR right trials, with a pre-movement increase in SR left trials and a post-movement increase in SR right trials. Because of the heterogeneity in the timing of responses across neurons and the different firing patterns in SR left and SR right trials, analyzing neural responses within fixed time intervals would not capture the full extent of this dynamic encoding, since fixed intervals assume uniformity in neural responses. Therefore, to capture the time course of changes in neural response for each neuron, we performed consecutive t-tests, combined with permutation analysis, between the firing rate during the baseline period (100ms before the onset of the stimulus) and the firing rate in a sliding window of 80ms, shifted in 10ms increments from the onset of the stimulus to 400ms after the response (see Methods). We defined the task-modulated neurons as those neurons whose activity was significantly different from the baseline activity at any time after the onset of the stimulus for at least 8 consecutive bins. Fewer neurons in EL, compared to OT and LL, showed a task modulated activity in both SR left (OT 70/80; 60/80 LL; EL 40/80, χ^2^ test, LL vs OT from stimulus p=0.01; χ^2^ test, EL vs OT from stimulus p=9x10^-11^; χ^2^ test, EL vs LL from stimulus p=5x10^-5^; χ^2^ test, LL vs OT from movement p=0.2; χ^2^ test, EL vs OT from movement 5x10^-9^; χ^2^ test, EL vs LL from movement 4x10^-6^), and SR right (OT 72/80; 61/80 LL; EL 47/80 ; χ^2^ test, LL vs OT from stimulus p=0.004; χ^2^ test, EL vs OT from stimulus 3x10^-14^; χ^2^ test, EL vs LL from stimulus 9x10^-7^; χ^2^ test, LL vs OT from movement p=0.1; χ^2^ test, EL vs OT from movement 3x10^-9^; χ^2^ test, EL vs LL from movement p=8x10^-6^) (Figures 2B,C).

The premovement response of individual neurons emerged at different times during the trial (Figure 2B). To find the precise moment when a neuron’s activity significantly diverged from the baseline, we identified the first instance within a sequence of at least 8 consecutive time bins where the activity was significantly greater than the baseline (p<0.05, see Methods statistical methods). Figure 2D illustrates the cumulative distribution of the proportion of neurons which showed a task modulated activity at any given time through the entire trial. Each point on the curve represents the proportion of neurons that started to respond within a specific time during the trial. When the activity was aligned to the onset of the stimulus, the cumulative distribution in EL was shifted to later time bins compared to LL and OT in SR left (Kolmogorov-Smirnov test OT vs EL, p=4x10^-4^; Kolmogorov-Smirnov test OT vs LL, p=0.3; Kolmogorov-Smirnov test EL vs LL p=0.003), but not in SR right trials (Kolmogorov-Smirnov test OT vs EL , p = 0.1; Kolmogorov-Smirnov test OT vs LL, p=0.9; Kolmogorov-Smirnov test EL vs LL p=0.2). However, when the activity was aligned to the onset of the movement, the empirical cumulative distribution in EL trials was significantly shifted to later time bins in both SR left (Kolmogorov-Smirnov test OT vs EL , p=0.02; Kolmogorov-Smirnov test OT vs LL, p=0.4; Kolmogorov-Smirnov test EL vs LL p=0.3) and SR right (Kolmogorov-Smirnov test OT vs EL , p=0.002; Kolmogorov-Smirnov test OT vs LL, p=0.7; Kolmogorov-Smirnov test EL vs LL p=0.05) trials.

**Figure 1.**
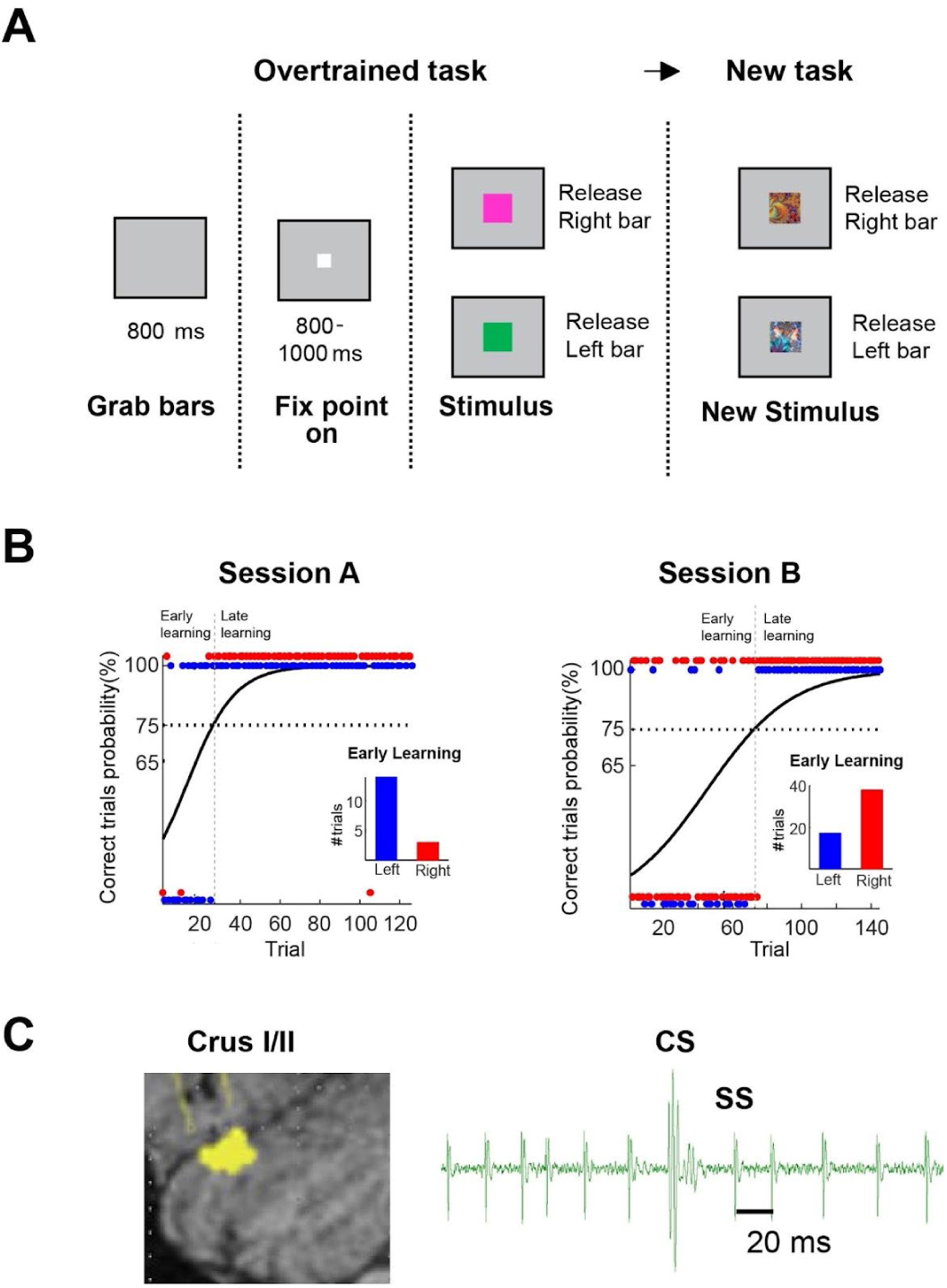
Behavioral task, Crus I and Crus II recording location. **A)** Visuomotor association task in overtrained and new task. A typical session started with the overtrained task. In each trial, a fixation point appears as soon as the monkey touches both the left and right bar. After a random time from 800 to 1000 ms, a stimulus appears. The green stimulus requires the release of the left bar, the pink stimulus requires the release of the right bar. The overtrained task is followed by the new task with two new stimuli that the monkey has never seen before. **B)** Binary logistic regression of the performance in two representative sessions. The y-axis is the probability that the trial on the x-axis is correct. Blue and red dots indicate left and right bar release, respectively. All trials with a probability of being correct less than 75% belong to the early learning; all trials with a probability higher than 75% belong to the late learning. Inset: number of trials for left (blue) and right (red) bar release in early learning, regardless of the trial outcome. **C)** Left: Area of Curs I and Crus II where recording was performed. Right: Simple Spikes (SS) and Complex Spike (CS) of a representative neuron.

**Figure 2.**
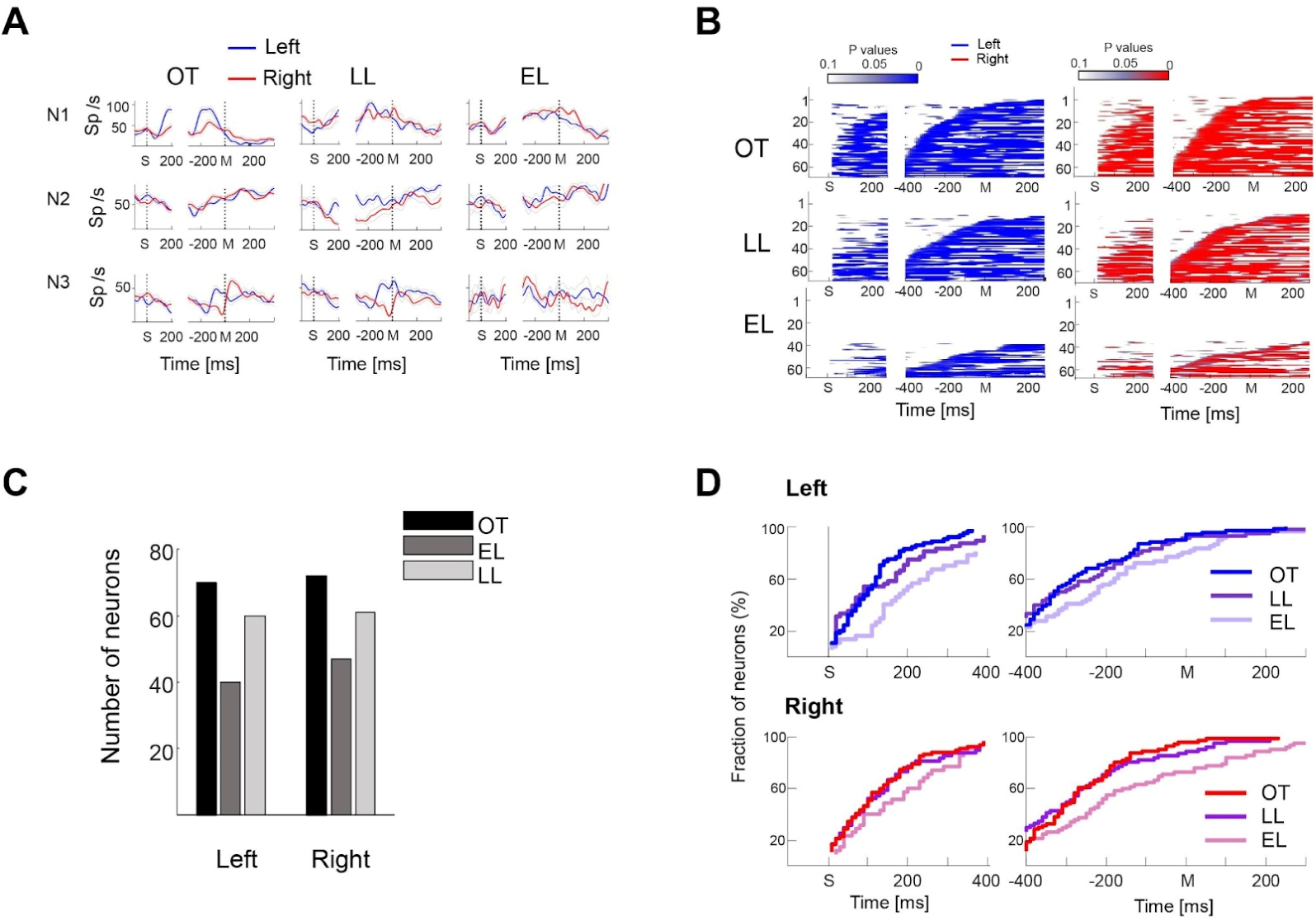
Purkinje Cells respond at different times during the trial to the Stimulus-Response association. **A)** Responses of three representative neurons in correct trials for left (blue) and right (red) response. OT: overtrained task; LL: late learning (new task); EL: early learning (new task). X-axis: times (ms) aligned with the onset of the stimulus (S) and from the onset of the movement (M). **B)** Task modulated neurons in 80ms time windows stepped by 10ms, after the stimulus onset and around movement onset, with respect to their baseline activity (100ms before the stimulus onset), for left and right correct trials (t-test). **C)** Total number of neurons in B) whose activity is significantly modulated for left and right correct trials in OT, EL and LL (p-value<0.05). **D)** Empirical cumulative distribution of the fraction of neurons in B) significantly modulated for left and right response in correct trials after the stimulus onset and around movement.

### Different Purkinje Cells discriminate between Stimulus-Response left and Stimulus-Response right at different times

Although all neurons responded to SR right and left in OT, the majority showed a SR selectivity. The activity patterns in EL were different from those in OT and LL. Moreover, the activity patterns in LL do not necessarily match the patterns in OT (see Neuron 2 in Figure 2A). These observations suggest a flexibility in neural response patterns, indicating that the response of PCells is not pre-determined, but can reorganize based on learning and task demands. When considering the neural population, the discrimination between SR left and SR right is not confined to a specific moment in time but extends from the stimulus onset into the period following the movement, regardless of the performance (Figure 3A). However, fewer neurons discriminate between the two choices during EL compared to LL and OT (Figure 3B), but as learning increased, more neurons became selective (Figure 3A left, 200ms after stimulus onset; χ^2^ test, EL vs OT, p=0.0006; χ^2^ test, LL vs OT, p=0.07 ; χ^2^ test, LL vs EL, p=0.1; Figure 3A right, 400ms before and 200ms after movement onset; χ^2^ test, EL vs OT, p=0.008; χ^2^ test, LL vs OT, p=0.4 ; χ^2^ test, LL vs EL, p=0.01). When considering the empirical cumulative distribution of the proportion of neurons that discriminate between the SR left and the SR right, the majority of neurons discriminate between the two associations earlier after the stimulus onset in OT and in LL compared to EL (Figure 3C left; Kolmogorov-Smirnov test EL vs OT p=0.02; Kolmogorov-Smirnov LL vs OT p=0.2; Kolmogorov-Smirnov test LL vs EL p=0.4) and earlier before the onset of the movement in OT and LL, compared to EL (Figure 3C right; Kolmogorov-Smirnov test t EL vs OT p=0.02; Kolmogorov-Smirnov test LL vs OT p=0.6; Kolmogorov-Smirnov test from stimulus LL vs EL p=0.007).

These results suggest that the difference in the timing of the upcoming choice between EL and LL is not related purely to the motor command but reflects the acquisition of the correct sensory-motor association. To test this hypothesis, we compared the empirical cumulative distribution of correct with error trials of the times when each neurons started to distinguish between SR left and SR right; this was only possible for the EL trials, because there were not enough errors in the LL trials. Similar to correct trials, in EL only a minority of neurons was able to distinguish between SR right and left in error trials during EL (Supplementary Figure 2A). In addition, like in correct trials, only a minority of neurons were SR selective in the error trials (Supplementary Figure 2A; χ^2^ test, from stimulus p=0.5; χ^2^ test, from movement p=0.7). Moreover, the empirical cumulative distribution of the times when the population started to discriminate between SR left and SR right in the error EL trials was not significantly different from that in the correct EL trials (Supplementary Figure 2B; Kolmogorov-Smirnov test, p=0.8 and p=0.4, aligned with the stimulus and with the movement, respectively), further highlighting a relatively low level of encoding during EL in general.

**Figure 3.**
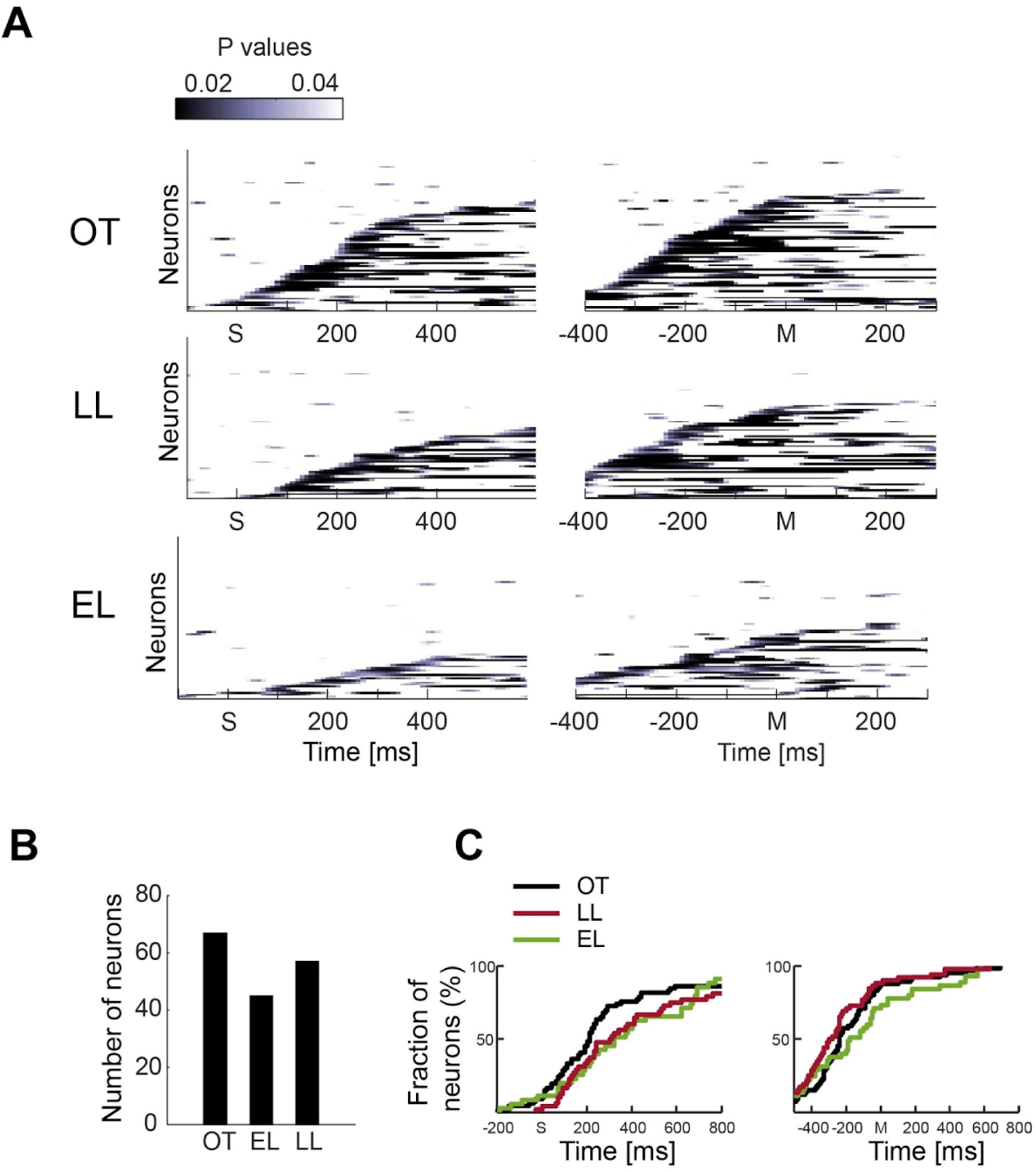
Different Purkinje Cells discriminate between Stimulus-Response left and Stimulus-Response right at different times. A) Response selective neurons after stimulus (S) and around movement (M) onsets (p value of t-test in a sliding window of 80ms), during overtrained (OT), early (EL), and late (LL) learning (new task). **B)** Number of response selective neurons (p-value<0.05) from A). **C)** Empirical cumulative distribution of the fraction of neurons in B) selective for the response.

### Stimulus-Response signal decoding

We decided to analyze further the signal dynamics and the stability of the signal throughout the trial at the population level. We disambiguated the pure visual signal from the response signal as a function of time in the OT, EL and LL stages of the visuomotor association learning task. We trained support vector machines^5,6^ with a linear kernel to decode from the neural activity of Purkinje cells the future response after the stimulus onset and before the movement onset, and the actual response after the movement onset. We analyzed correct and error trials separately. In this first analysis we could not disentangle the response signal, i.e., right and left, from the stimulus identity signal, i.e., stimulus 1 and stimulus 2, and we will continue referring to this signal as to the stimulus-response (SR) signal. We trained and tested linear decoders on pseudo-simultaneous trials (pseudo trials). We defined the pseudo trial as the combination of spike counts randomly sampled from different trials of the same task condition. We computed, for each neuron, the z-scored activity in a 200ms time bin, stepped by 20ms. In order to have enough training and testing trials for the linear decoders, we analyzed the activity of neurons recorded for at least 5 correct trials in each response condition (right and left) in OT, EL, and LL for a total of 54/80 neurons in OT, 18/80 neurons in EL, and 42/80 neurons in LL. The decoder identified the monkey’s response after stimulus onset (Figure 4A), and before movement onset (Figure 4B) for correct trials. The average reaction times (mean±SEM) in OT, EL, and LL, are (412±4)ms, (760±24)ms, and (519±5)ms, respectively (Supplementary Figure 3).

We subsequently analyzed error trials during the novel task (NT), and we retained those sessions with at least 5 trials per condition, for a total of 24/80 neurons. The analysis of error trials showed that the SR is not decoded immediately after the stimulus onset. Instead, it occurs only after approximately 600ms, which is the average reaction time during error trials (Figures 4C-D). The signal dynamics during error trials in EL were similar to the dynamics of correct trials in EL, where there was no significant signal after the stimulus onset (as seen in Figure 4A, red curve). In error trials, around the movement onset, the signal became significant only at the moment of the response, but it dropped to chance in ∼300ms after the response. This was different from the dynamics of correct trials in NT where the signals remained significant for ∼400ms in EL and ∼600ms in LL. The average reaction time in error trials was (691±17)ms. We observed high variability in the reaction times across trials and across sessions, particularly in correct trials in EL and error trials (see Supplementary Figure 3 for full distributions).

We also decoded the outcome of the previous trials before the stimulus onset and the outcome of the current trial around the movement onset during early learning. We found that the previous outcome was significantly encoded in the 500ms preceding the stimulus onset (Figure 5A). The outcome of the current trial was significantly decoded ∼400ms after the movement onset (Figure 5B).

**Figure 4.**
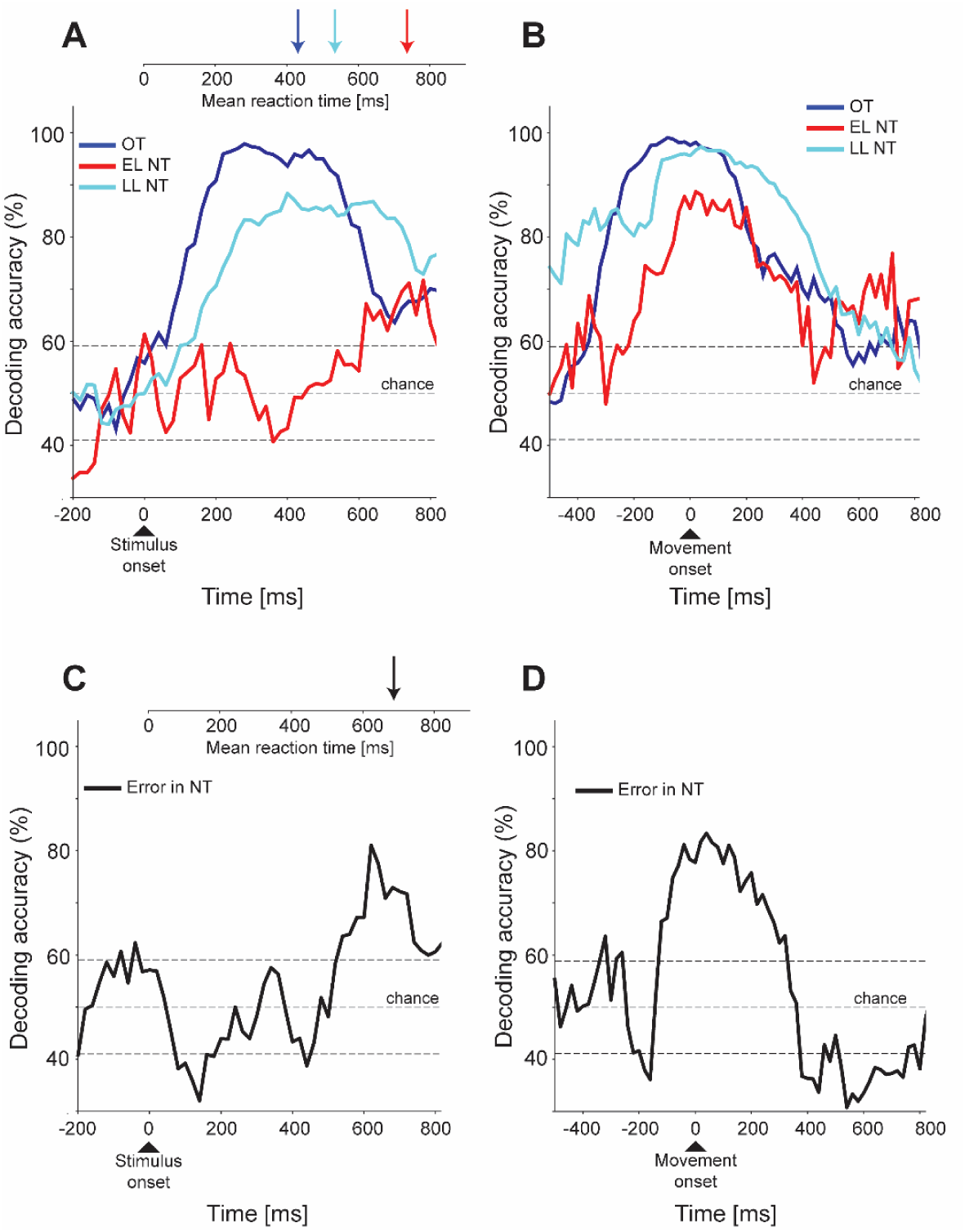
Population linear decoding of the Stimulus Response (SR) signal during Overtrained task (OT), in Early (EL) and Late Learning (LL) of Novel Task (NT). **A)** Performance of a linear classifier in decoding the response around stimulus onset in OT (blue curve), in the EL (red curve), and LL (cyan curve) for correct trials. On top of the decoding accuracy, the vertical arrows indicate the mean reaction time. The horizontal-dashed lines are ±2 standard deviations of the distribution of decoding accuracies from 100 null models with random shuffles of the labels. **B)** The same as in A) but around the movement onset. **C)** Performance of a linear classifier in decoding the response around stimulus in error trials during NT. On top of decoding accuracy, the vertical arrows indicate the mean reaction time. The horizontal-dashed lines are ±2 standard deviations of the distribution of decoding accuracies from 100 null models with random shuffles of the labels. **D)** The same as in C) around the movement onset.

**Figure 5.**
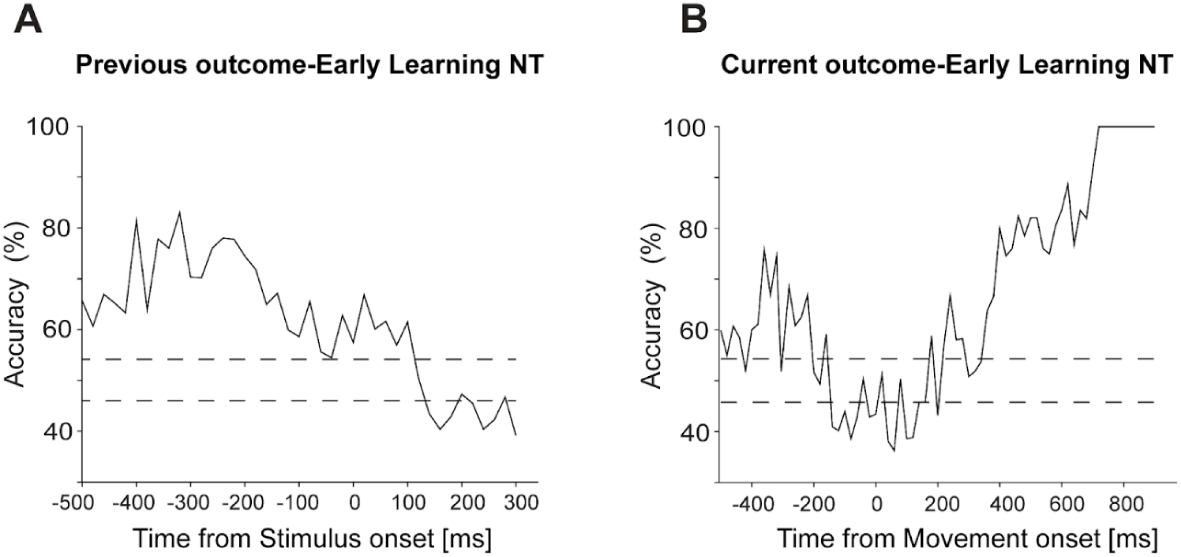
Decoding of previous and current outcome. **A)** Decoding of the outcome of previous trials around current stimulus onset during Early Learning of Novel Task (NT). **B)** Decoding of the current outcome around the movement onset during Early Learning of Novel Task (NT).

### Stimulus-Response signal stability analysis

To test the stability of the signal over time, we used cross-bin decoding, which involved training a linear decoder in one time bin and testing it in a different time bin. If the decoder’s accuracy in generalizing across time bins differs from the chance interval obtained from the distribution of accuracies of null models, it suggests that the variable’s representation is consistent over time. Figure 6 shows the cross-bin decoding accuracy of the linear decoder in classifying the response during OT and NT. For this analysis, we combined the data from EL and LL (see Supplementary Figure 4 for the results separately for EL and LL). We found that the response representation is more stable during OT than NT after the stimulus onset (as shown in Figures 6A,B). We defined the time interval during which the generalization accuracy is significant before it reaches chance level as the difference between the time bin used during training and the first testing time bin where the decoding accuracy is no longer significant. We only did the analysis in the forward direction, where the testing time bin is the same as or after the training time bin. After the stimulus onset, the time interval with a significant decoding accuracy is longer in OT (295±34ms, mean ±SD) than in NT (154±37 ms) (Supplementary Figure 5A; Mann-Whitney U test; p-value=1.2^-11^). This finding suggests that after the stimulus onset, if the information is significantly encoded in both tasks, the response representation is less stable across time in NT than OT, indicating a faster reconfiguration of the decoding weights (Supplementary Figure 6). Interestingly, after the movement onset, we observed that the signal is more stable across time in NT (187±106ms) than OT (140±74ms) (Mann-Whitney U test; p-value = 0.001, see Supplementary Figure 5B).

**Figure 6.**
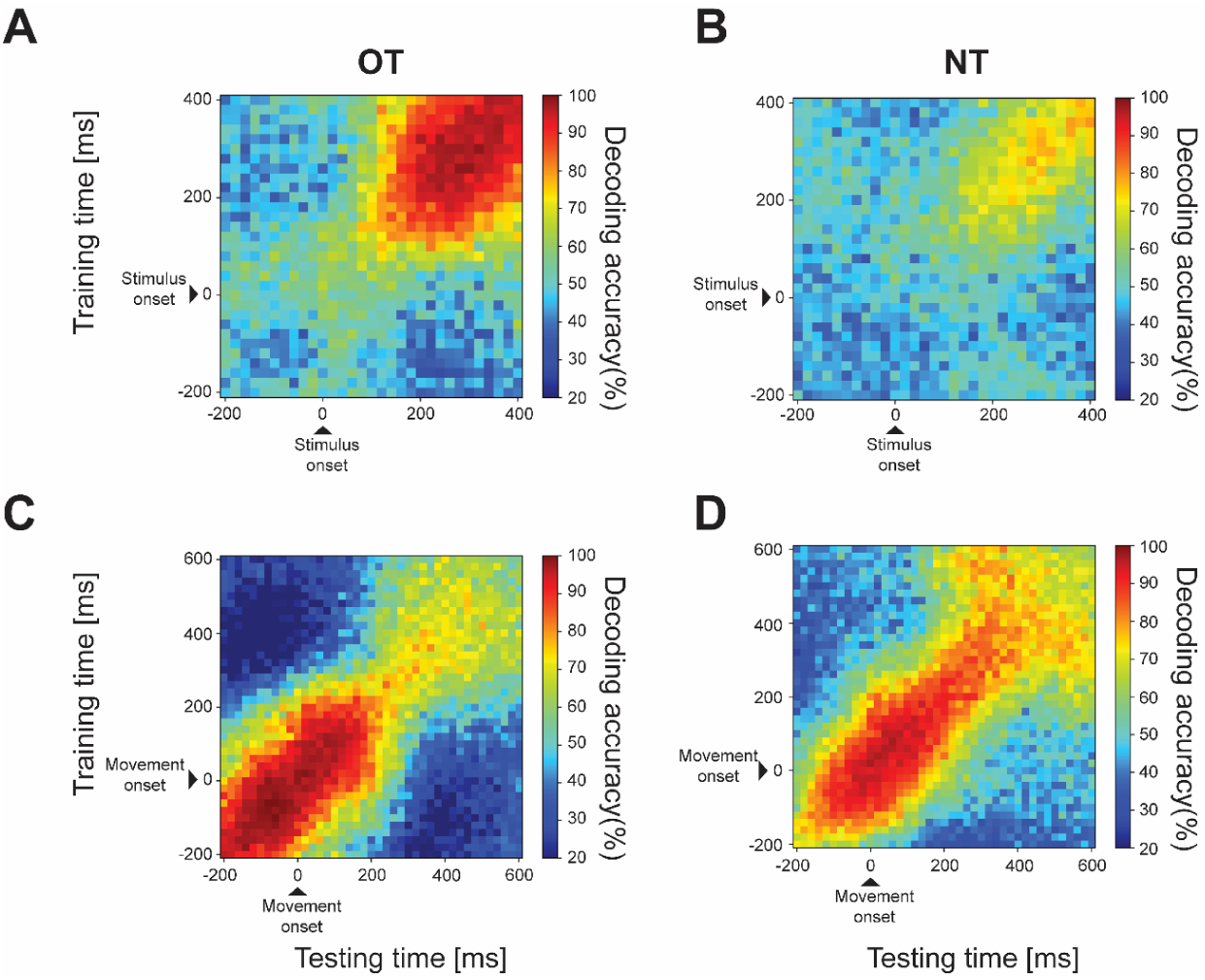
Cross bin decoding in correct trials during Overtrained (OT) and Novel (NT) tasks. **A)** Cross bin decoding around stimulus onset during OT. The linear decoder is trained on a time bin (y-axis) and tested on other time bins (x-axis) preceding and following the training bin. The diagonal contains the decoding accuracy where the decoder is trained and tested on the same time bin. The color bar is the performance of the linear decoder in generalizing across time bins. The same as in A) during NT. **C) C**ross bin decoding around movement onset during OT. **D)** Same as in C) during NT.

### Neural decoding of the response and stimulus identity signals

To differentiate the amount of information encoded by the Purkinje cells for the response (right and left) from the visual signal (stimulus 1 and stimulus 2), we examined the correct trials of the novel and reversal tasks together. In the reversal task (REV), the visual stimulus and the response are reversed with respect to the novel task, making the response and stimulus identity uncorrelated. We defined the early and late learning phases as those trials in which the average behavioral performance in each session is between 40% and 60% and above 70%, respectively, in both novel and reversal tasks. We chose these thresholds to maximize the number of trials and neurons both in NT and REV tasks. During EL, the decoder could not identify the stimulus identity after the stimulus onset, except for a small bump after ∼200ms from the stimulus onset (Figure 7A). During LL, the signals’ dynamics change. The decoder identifies stimulus identity after ∼100ms from the stimulus onset, followed in time by the response signal (Figure 7C). In EL, around the movement onset, the decoder identifies the stimulus identity before the movement onset (Figure 7B). This signal is maintained after the movement onset for ∼200ms, unlike in LL, where the stimulus identity signal drops to chance right after the movement onset (Figure 7D).

We also decoded the response and stimulus identity signals after combining the correct and error trials during EL and LL. Supplementary Figure 7A shows that, during early learning, there were no significant signals above chance in the 200ms following EL, LL, and REV. However, during LL, the stimulus identity is decoded shortly after the stimulus onset, followed by the response signal (Supplementary Figure 7C). This difference might be explained by the nature of the tasks being analyzed. Indeed, during the reversal task, the animals were not new to the stimulus identity, as they had previously been trained on the same stimuli in the novel task. In the reversal task, they were asked to reverse the stimulus-response association with respect to the association learned in the novel task. In contrast, during the novel task, the stimulus identities were entirely new to the animals, which is why there was no significant signal after stimulus onset in the early and late learning sessions (Supplementary Figures 7A,C). This differs from what was observed when we analyzed correct trials in the novel and reversal tasks (Figures 7A,C).

**Figure 7.**
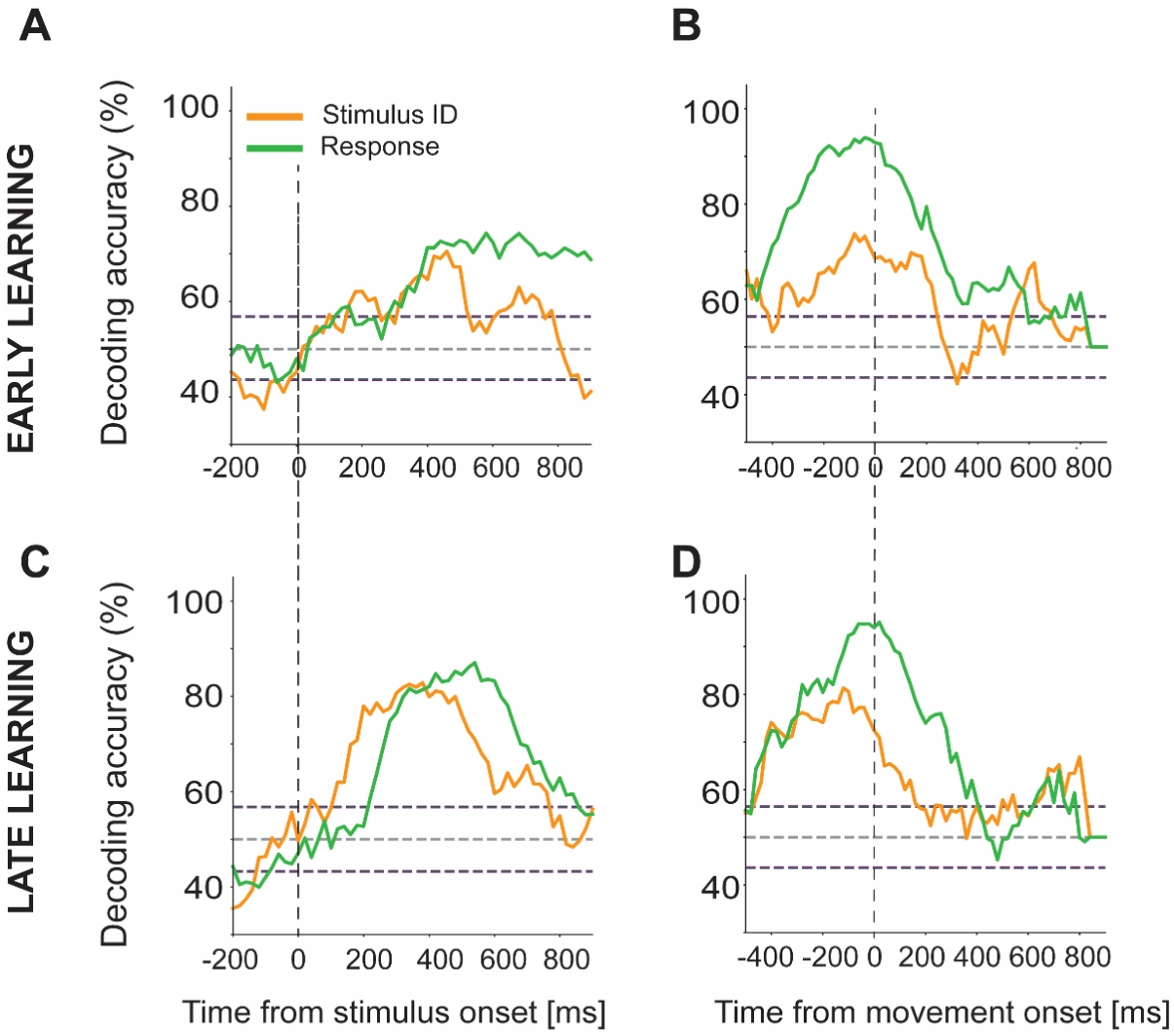
Decoding of the response and stimulus identity in the Early and Late Learning phases of Novel (NT) and Reversal (REV) tasks using correct trials. **A)** Stimulus identity (orange curve) and response (green curve) signal from the stimulus onset after combining the correct trials in the Early learning of NT and REV task. **B)** The same as in A) around the movement onset. **C)** Stimulus identity (orange curve) and response (green curve) signal after the stimulus onset after combining the correct trials in the Early learning of NT and REV task. It is evident that the stimulus identity is decoded after ∼100ms from the stimulus onset, followed in time by the response signal. **D)** The same as in C) around the movement onset. The horizontal dashed lines are 2 standard deviations of the distribution of decoding accuracies obtained from 100 shuffles of the labels.

### Potential mechanism underlying different timing of EL and LL signals

To investigate the potential mechanism behind the different timing of stimulus and response signals in the EL and LL, we conducted two modeling studies aimed at reproducing the neuronal signals depicted in Figure 7. In the first model, we hypothesized that the visual input to the PCs became more specific during learning. In the second model, we trained a simple feedforward neural network to perform the NT and reversal (REV) tasks. After achieving high accuracy in learning the tasks, we analyzed the activation of the artificial units over time, employing the same analyses that we applied to real PCells activity.

### Model of the dynamics of the input to Purkinje Cells

Our aim was to replicate the neural signal dynamics of PCells in both the EL and LL after the stimulus onset illustrated in Figures 7A,C. Indeed, during EL the stimulus signal was decoded after ∼300ms from the onset of the stimulus (with a brief but significant bump between 200ms and 300ms) almost simultaneously with the response signal (Figure 7A). On the other hand, when the monkeys learned the new association, the dynamics of the signals became clearer. The visual signal was decoded after ∼100ms from the stimulus onset, followed in time by the response signal which was significant after approximately 200ms from stimulus onset (Figure 7C).

We hypothesized that during the early learning stage, the visual input signal to PCells was weak and below the activation threshold of PCells (Figures 8A-C). This made it difficult for the visual input to be decoded by itself without the response input signal, which was above the threshold (Figure 8B, left). However, in the late learning stage, once the visuomotor association was well established, PCells might have received the visual signal above threshold earlier than the response signal, and we could decode the visual signal before the response signal (Figure 8B, right). To test this hypothesis, we developed a model consisting of 200 artificial input units. Out of these units, 50 units each encoded the visual stimulus 1 and 2, and the remaining units encoded response 1 and 2. The input signals were randomly projected to 100 output units with the Rectified Linear function as the non-linear activation function (Figure 8A). We decoded the visual and response signals from the activity of the output units, as we did with the neural data. The results showed that during early learning, the visual signal was barely decoded after the stimulus onset, and it became decodable along with the response input (Figure 8C, left). On the other hand, when we provided a stronger visual input above the activation threshold followed in time by the response signal, we observed a different output signal dynamic that resembled the neural signal dynamics observed in the late learning stage of actual data (Figure 8C, right).

**Figure 8.**
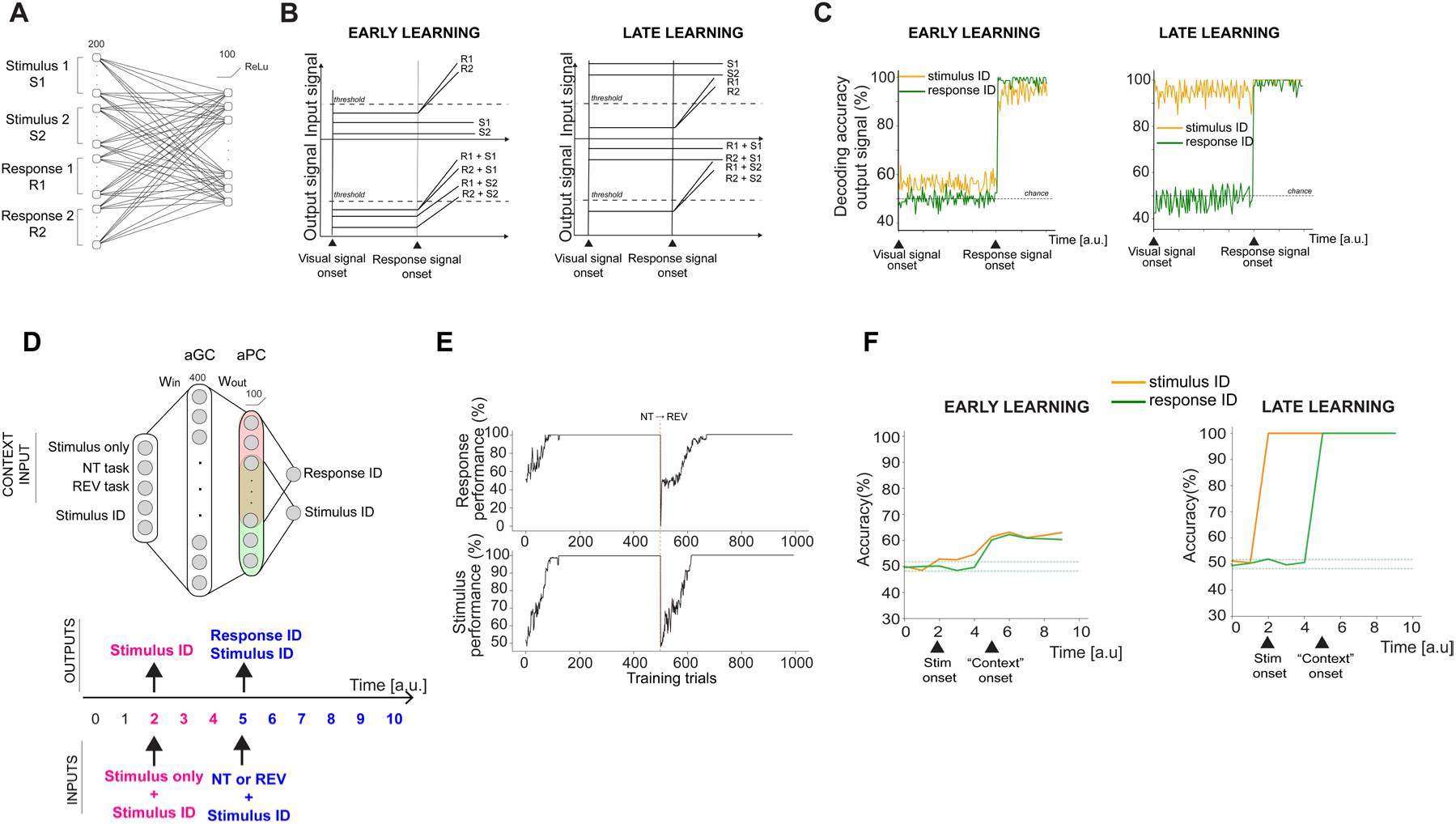
Schema of the input dynamics signal to Purkinje cells and architecture, performance, and decoding results of the neural network trained to perform the new and reversal tasks. **A)** Schematic of 200 input units conveying the visual (Stimulus ID) and response (Response ID) inputs. The input units are fully connected to 100 output units. The weights are randomly drawn from a uniform distribution between 0 and 1. The output current from input units is passed through a non-linearity defined by the rectified linear unit (ReLU). **B)** Schema of the input (top) and output (bottom) signal during early and late learning. During the early learning, the visual signal is below the activation threshold, and it is not decodable from the output activity. **C)** Decoding accuracy for stimulus and response signal in early and late learning given the input signals modeled as in B). **D)** Top: Architecture of the feedforward neural network: 5 input units with a one-shot vector of 3 units encoding the task context, i.e. stimulus only, NT, or REV, or stimulus only, and another one-shot vector of 2 units encoding the stimulus identity The input is projected to an expansion layer of 400 units (aGC) through random connection weights (Win) randomly initialized from a uniform gaussian distribution with positive values. These input weights are not trained. The output from the aGCs is passed to 100 units layer (aPC). Two readout units are trained to provide the correct output response and stimulus identity by reading the output activity from two subgroups of aPC (red and green areas) with an overlap of 20% (yellowish area). The network is trained to provide the correct response associated with the input stimulus and context, and the stimulus ID. Bottom: schematic of the dynamics of the inputs along a trial. **E)** Performance of the network during 1000 training trials. At 500^th^ trial there is the task switch, from NT to REV task. **F)** Decoding accuracy of stimulus and response across time from the aPC activity output during early and late learning. The horizontal dashed lines are 2 standard deviation from the mean of 100 null models. ReLU: Rectified Linear Unit; aGC: artificial Granule Cell; aPC: artificial Purkinje Cell; NT: Novel Task; REV: reversal task.

### Training of an artificial neural network to perform the new and reversal tasks

We trained a simple feedforward artificial neural network (ANN) to perform the NT and REV tasks. The ANN had 5 input units, which included a one-hot vector of 3 units encoding if it were a stimulus-only, the NT, or REV context. The other two units encoded the identity of the stimulus, (Figure 8D). The reason for the stimulus-only context input was to reproduce the dynamics of the visual signal prior to the response signal after stimulus onset. The NT and REV context inputs were necessary because the simple ANN could not achieve high performance on the reversed association, so we were forced to cue the task switch, even though it was not true for the real experiment.

The input was passed through an expansion layer of 400 units (which we referred to as artificial Granule Cells (aGCs)) for a more biologically plausible interpretation of the model) through random connection weights W_in_, which were randomly initialized from a uniform Gaussian distribution with positive values. These input weights were fixed throughout the whole training. The output from the aGCs was passed to a 100 units layer which we labeled as the artificial Purkinje cells (aPCs) layer. The output signal from aGCs was sent to aPCs through W_out_ weights that have been trained using a specific learning rule (see Methods). The network was trained such that an output unit had to provide a binary response (Response ID) associated with the input stimulus and context reading out the activity of 60 aPCs; a second output unit had to provide the visual stimulus identity (Stimulus ID) from the activity of other 60 aPCs, with an overlap of 20% with aPCs units readout from the Response ID unit. This is done in order to disambiguate the stimulus input signal from the output response signal in the aPCs activity (Figure 8D). We trained the ANN to perform the NT and REV tasks in two separate blocks of 500 trials each, for a total of 1000 training trials. In the first 500 trials, the network was trained to receive the stimulus-only and the NT contexts along with a Stimulus ID randomly chosen in each trial NT task, while in the last 500 trials it was trained to learn the reversed stimulus-response association (REV task). The dynamics of the input’s signals were the same as the ones modeled in Figure 8B, with the difference that now we did not impose any distinction in the input dynamics in EL and LL. In particular, we first provided the stimulus-only context and stimulus ID for a few time steps, and then we provided the NT or REV context along with the stimulus ID for the following time bins (Figure 8D, bottom). As before, we modeled a PCell’s activity as non-linear function of the inputs (ReLu). Both stimulus ID and context were randomly chosen at the beginning of each trial. In the case of the stimulus-only input, the network was required to give the stimulus identity as output. In the case of EL and LL, given a random input stimulus and context, the network had to learn the correct response associated to the input stimulus within the input context. After 500 trials, we reversed the stimulus-response association, and we randomly re-initialized 5% of the W_out_ weights of aPCs units read out only from the Stimulus ID output unit (see Figure 8D), so the network had to learn again to both encode the stimulus-only input and reversed association. Figure 8E shows the network’s performance in classifying the input stimulus (top row) and the correct response associated to the input stimulus (bottom row) along the training trials. We defined, as done for the real experiment, an early learning stage as the first trials of NT and REV where the performance of the ANN for the correct response was below 60%. We defined the late learning stage the last 100 trials of NT and REV, where the ANN performance for correctly classifying the response was 100% (Figure 8E).

We decoded the stimulus and the correct response from the output signals of the aPCs using the same analysis tool used on actual data. Figure 8F shows the decoding accuracy of the response (green line) and stimulus (orange line): during EL, at time step 2, when the Stimulus ID was provided to the ANN, we could not decode it from the aPCs activity, but only at time step 5 with the response (Figure 8F, left). This dynamic resembles the neural results we observed in the early learning stage of the monkey, where the stimulus identity was decodable only along with the response signal (Figure 7A). This suggested that, even though the stimulus was provided in the inputs, it was not decodable from the aPCs because it was below the activation threshold (see schematic in Figure 8B, left). Differently, in the late learning phase of the ANN where the performance was 100%, the visual stimulus identity was decodable as soon as it was provided as input to the network, followed in time by the response signal (Figure 8F, right). This dynamic resembled the dynamics of the neural signal observed in the late learning stage, where the visuo-motor association was well established (Figure 7C). This indicated that when the ANN learned the stimulus-response association with high accuracy, the output signal for the stimulus identity was above the aPCs activation threshold and then significantly decodable prior to the onset of the response, as in the actual data.

## DISCUSSION

When monkeys perform a visuomotor association task, neurons in the posterolateral cerebellum respond in the interval between stimulus appearance and response. The response begins at different times in the trial for different neurons, with identification times starting close to the stimulus onset for some neurons and continuing until the response. More neurons respond during the overtrained task (OT) and late learning (LL) epochs than during the early learning (EL) epochs. The identification times occur closer to stimulus appearance and longer before the response in OT and LL than in EL. Although all neurons that responded before the movement of one hand also responded before the movement of the other hand, a majority of the neurons had a significant difference in the intensity of their response between left- and right-hand movements. This difference also appeared earlier after the stimulus and longer before the movement in OT and LL than in EL. A linear decoder identified not only the time at which information about movement occurred and the hand used but also identified the identity of the stimuli that ordered the monkey’s response.

Given this clear correlation between neural activity and behavior, it is tempting to assume that this activity is important in driving the monkey’s decision. However, because inactivation of the posterolateral cerebellum with muscimol has no effect on the numerical performance in OT^3^, during which the correlation between neural activity and behavior was strongest, the Purkinje cell activity in the crus regions cannot be essential for driving the decision process. At best, their SS responses may contribute to the early learning process and to accelerating the ultimate visuomotor response, as muscimol inactivation impairs their response during learning. Thus, decisions can be made well without the Purkinje cells in the posterolateral cerebellum, once the decision process has been acquired.

The pattern of movement responses in the visuomotor association task resembles that seen in monkey prefrontal cortex during a visuomotor association reversal task^7^. In the saccade task, unlike in our visuomovement association task, neurons only fire before saccades of one direction, as opposed to the neurons described here, which fire before movements of both hands, although with differing intensity. The immediate presaccadic response occurs before every saccade, correct or error, but only begins to approach the time of stimulus onset as the monkey learns the correct saccade direction^8,9^. Given the resemblance in the relationship of neural activity to movements in both prefrontal cortex and posterolateral cerebellum, and the anatomical relationship between prefrontal cortex and posterolateral cerebellum, it is likely that the cerebellar movement activity is a corollary discharge arising in prefrontal cortex and/or the basal ganglia. This suggests that the prefrontal cortex and basal ganglia are actually driving the decision process, and that they may engage the cerebellum for facilitating the learning process, accelerating execution of the decision process and rendering the decision process more efficient. This concept is compatible with studies that investigated less complex discrimination behaviors in mice^10,11^.

In order to create the error signal that facilitates visuomotor association learning the cerebellum needs both information about the outcome of the prior trial, the prior movement, and the current stimulus. This information is present in the activity of the SS. In most forms of motor learning, the complex spike signals, which all reflect climbing fiber activity originating in the inferior olive, can provide such an error signal^12–15^. However, the complex spike activity in our visuomotor association task does not provide an obvious error signal^16^. Instead, when the complex spikes responses are classified based on the SS properties of the Purkinje cells, they appear to encode the probability of failure in the current trial, but not the actual results of the prior trial. In addition, they report the cue signaling at the beginning of the trial after stimulus onset as well as information that correlates with the context and learning state. The SSs begin to fire, and identify the hand used, at different times during the trial. This tiling of the interval from stimulus appearance to response by the population of neurons resembles the tiling of reward-based activity in SSs, which has also been observed during our visuomotor association task^2^. Similar tiling has also been observed in the granule cell layer of lobule simplex in relation to learning-dependent timing of eyeblink responses^17,18^, and it also occurs in the striatum to process information at longer time scales^19^. The utility of tiling is not well understood. It may save energy if the activity can be summed by recipient neurons, such as the cerebellar nuclei neurons, which integrate signals from about 40 Purkinje cells^20,21^. Alternatively, in the motor system it may be related to the stereotypical temporal recruitment of motor neurons during increasing intensities of force production^22^.

Our computational results show that the identity of the stimulus, not found in the activity of single neurons, can be found in the activity of the population taken as a whole. Recent evidence in mice and in non-human primates indicates that multimodal information processing can occur to a large extent through the collective activity of Purkinje cells^23–33^. The complex microcircuits of the cerebellar cortex, with recurrent feedback and the combination of different forms of plasticity during learning, may account for the different latencies of the premovement responses of different Purkinje cells we reported in this study^34^. Context-dependent signals begin at the level of granule cells^17^, which integrate multimodal convergent and divergent inputs from mossy fibers^35,36^. Over a hundred thousand granule cell axons provide a single Purkinje cell with contextual information in a simple association task^17,37,38^. The axons of the Purkinje cells establish inhibitory synapses to a single neuron of the cerebellar nuclei^20,21^, and, through their collaterals, with granule cells^39^ and molecular layer interneurons^40^. The collateralizations to the granule cells and to the molecular layer interneurons provides a mechanism of recurrent internal feedback which, in combination with different forms of plasticity^41,42^, may play a role in maintaining the balance of excitatory and inhibitory signals within the cerebellar cortex^18,27^ and allow the Purkinje cells to adapt to new information and changing task demands through the dynamic reconfiguration of neural circuits.

In conclusion, the different firing rate patterns we observed may reflect that different populations of Purkinje cells within the cerebellum may be engaged in parallel processing of different aspects of the task, potentially contributing to the overall computation of the sensory motor transformation during the decision-making process, and the diffuse synaptic plasticity within the circuits may provide flexibility and adaptability in the face of changing task demands. The finding that different neurons exhibit task modulation at distinct temporal windows suggests that the encoding of task-related information occurs in a temporally diverse manner. This temporal diversity may reflect the parallel processing nature of the cerebellar circuitry, with different populations of Purkinje cells engaged at different stages of the task. By encoding task information at different time points throughout the trial, different Purkinje cells may participate in the integration and coordination of sensory input, motor planning, and execution. The observation of a gradual shift in response latency may provide a process of circuit rearrangement and the establishment of new, task-specific neural pathways. Furthermore, the observation that simple spikes do not simply revert to a predefined state during learning^43^, implies that the cerebellar circuitry forms novel connections that reflect the specific demands of the new task, allowing the cerebellum to adapt to new information and changing task demands through the dynamic reconfiguration of neural circuits.

## MATERIALS AND METHODS

### Behavioral tasks

The monkeys were seated in a Crist primate chair. positioned inside a dimly lit recording booth with their heads securely immobilized. The visual stimuli, generated by Hitachi CP-X275 LCD projector, were presented to the monkeys on a back-projection screen positioned in front of the monkeys, at a distance of 57 cm. A Dell PC running the NIH VEX graphic system controlled the Hitachi CP-X275 LCD projector. We used the NIH REX system for behavioral control (Hays et al., 1982). Each trial began with the monkey grasping two bars with the left and the right hands. Following this, the trial initiation cue (a 1° x 1° white square) was presented for 800ms. Subsequently, one of the two stimuli appeared either briefly for 100ms or remained visible until the monkey initiated a hand response by releasing one of the bars. The stimulus’ presentation indicated which hand the monkey should release: one stimulus for the left hand and another for the right. A correct response—releasing the hand associated with the presented stimulus—was immediately rewarded with a drop of juice, accompanied by an auditory beep signaling the release of the reward. The reward and signal were delivered with a delay of 1ms following the initiation of the correct hand movement. From the initiation of the hand movement, a 700 ms delay (inter-trial interval, ITI) was enforced before the commencement of the next trial. Trials where both hands were released were aborted automatically.

Monkeys were trained to associate the specific pair of stimuli (a green and a pink square) with the corresponding hand releases over a period of 4-6 months until they achieved a performance accuracy above 95% (the overtrained, OT, condition). Each recording session began with trials in the overtrained condition to ensure consistent performance before introducing novel stimuli.

After a set number of trials in the overtrained condition, the stimulus pair was switched to two novel fractal images, previously unseen by the monkeys and not color-matched to the overtrained symbols. Over approximately 50-70 trials, monkeys demonstrated learning by improving their performance in associating each new stimulus with the correct hand release.

### Data Analysis of behavioral performance

To quantitatively assess the learning curve in response to novel conditions, we employed a binary logistic regression model with the binary outcome being the correctness of the trial (1 for correct, 0 for error). We evaluated the goodness of fit of each a binomial logistic regression model using a χ^2^-statistic vs. constant model. This model allows us to estimate how the probability of a correct response changes over the course of the learning period as the monkeys become familiar with the novel stimuli.

#### Quantification and Statistical Analysis

##### Single unit recording

For daily recordings, we utilized glass-coated tungsten electrodes with an impedance range of 0.8-1.2 MOhms (FHC, Inc.). These electrodes were introduced into the target cerebellar region using a Hitachi microdrive system. The raw signal captured by the electrode was initially processed through a Neurocraft head stage and an amplifier provided by FHC, Inc. To isolate the frequencies of interest, the signal underwent bandpass filtering using a Krohn-Hite filter, with a lowpass setting at 300 Hz and a highpass setting at 10 kHz, employing a Butterworth filter configuration. The filtered signal was then digitized and processed through a Micro 1401 system (CED electronics). The neural data acquisition was integrated with the NEI REX-VEX system, coupled with the Spike2 platform, to ensure synchronized collection of event-related and neural activity data. This setup allowed for real-time monitoring of neural signals and task events, crucial for aligning neuronal responses with specific behavioral outcomes. In both online (during recordings) and offline (post-experiment) analyses, we focused on identifying cerebellar Purkinje cells, characterized by their distinct complex and simple spike activity patterns. The identification process involved analyzing the spike waveform characteristics to confirm the isolation of Purkinje cells and to ensure the stability of recorded waveforms throughout each experimental session. A representative recording demonstrating simple and complex spike patterns is illustrated in Figure 1C (rightmost panel). To quantify the neuronal activity, we applied a spike density function generated using a Gaussian kernel with a standard deviation (sigma) of 15 milliseconds.

#### Single neurons analysis

##### Identification of hand response

To investigate whether neurons exhibit a task modulation in both SR left and SR right trials we conducted analyses on neuronal responses when monkeys executed these movements correctly. We first divided the responses of each neuron based on correct SR left -hand and SR right trials including only neurons with at least 4 trials for each group. Spike density functions were computed for each trial, averaging the response within 80ms epoch bins with a 10ms shift from stimulus onset and movement onset. We compared the mean activity of these bins with baseline activity, defined as the activity in the 100ms period preceding stimulus onset. Normality tests were performed for each epoch bin, followed by either a t-test for normally distributed data or a Wilcoxon rank-sum test for non-normally distributed data. If the p value was lower or equal than 0.05, we repeated the statistical tests (t-test or Wilcoxon test) after shuffling the labels of the baseline and of the responses 100 times. and calculated the fraction of the p-value was less than or equal to 0.05. We considered the difference for each bin significant when both the original t-test was significant and the fraction of significant p-values from permutation testing was less than or equal to 0.05. Divergence from baseline was identified as the first occurrence of at least 8 consecutive significant bins. Additionally, we repeated the analysis using spike counts instead of spike density functions, yielding consistent results across both methods. To identify the temporal dynamic across the population of the times when the response to the left and to the right-hand correct trials emerged, we performed an empirical cumulative distribution of the times when the difference between the neurons response and the baseline emerged across the neurons. To evaluate the differences between two distributions we performed a Kolmogorov-Smirnov test.

##### Identification of hand preference

All the statistical analysis, normality test, t-tests, Wolcoxon test, and Kolmogorov-Smirnov test have been performed using Matlab. To identify if a neuron responded differently in SR left and SR right trials, we performed, for each trial, a spike density function. We then computed the difference between at least 4 trials in in left hand response and at least 4 trials in right hand responses, in consecutive 80ms bins with a 10ms shift, within two time windows: a 400ms window starting from the onset of the stimulus and a 800ms window starting 400ms before the movement. For each bin we tested the distribution of each group for normality. If both distributions were normal, we compared the two-distribution using a parametric test (t-test). If not, we used a non-parametric test. If the p values were < 0.05, we pooled the SR left and the SR right hand trials, randomly selected two groups with the same number of trials as the condition with the minimum trials, and then performed the statistical tests (t-test or Wilcoxon test) for each bin.

We repeated the statistical tests (t-test or Wilcoxon test) after shuffling the labels of the pooled trials 100 times. and calculated the fraction of the p-value was less than or equal to 0.05. We considered the difference for each bin significant when the original t-test was significant and the fraction of significant p-values from permutation testing was less than or equal to 0.05. We repeated the analysis using spike counts instead of spike density functions, yielding consistent results across both methods.

To evaluate the differences between two ECD distributions we performed a Kolmogorov-Smirnov test. To confirm that the observed differences between two distributions were not the result of random variations, we randomly reassigned trial identities within the dataset and reanalyzed the data for each permutation. The permutation was repeated 100 times.

The significance of the differences was determined based on the proportion of permutation outcomes that produced a p-value less than 0.05. We consider a significant difference between two distributions of the p-value of the original results less than 0.05, and the proportion of significant p-value of the permutation was less than 0.05.

#### Neuronal population analysis

##### Neurons and trials selection, pseudo trials definition, and decoding of neuronal population activity

We analyzed the neuronal activity recorded in the overtrained (OT), novel (NT) and reversal (REV) tasks. During NT and REV tasks, we defined each session’s early and late learning phases as those trials with an average performance below and above 75%, respectively. We subsequently analyzed the neuronal activity of correct trials in early and late learning separately. In particular, we analyzed those sessions with at least 5 trials per condition, i.e., right and left responses, in correct and error trials. We analyzed error trials only in early learning because there were not enough trials per condition during the late learning phase.

Linear decoders were trained and tested on pseudo-simultaneous population trials (pseudo trials). We defined a pseudo trial as the combination of spike counts randomly sampled from every neuron in a specific time bin and task condition ^44^ . Pseudo trials were generated as follows: given a time bin *t* and task condition *p*, for every neuron, we randomly picked a trial of task condition *p*, and we computed the spike count in the time bin *t*. The single pseudo trial ξ, for condition *p* at time bin *t*, is then *p*(t)=(ξ_1_^p^(t), ξ_2_^p^ (t),…, ξ_Ν_^p^ (t)), where *N* is the number of recorded neurons, and ξ_i_^p^ (*i* is the neuron identity, *i*=1,…,N) is the spike count. We repeated this procedure 100 times, with 1000 pseudo trials per task condition and time bin. To disambiguate the visual stimulus from the response signals, we analyzed correct trials in the NT and REV tasks together, in the early and late phase, separately.

For each binary variable, i.e., stimulus and response, we trained a Support Vector Machine (SVM) classifier with a linear kernel to classify the spike count into either of the two values of the binary variables– stimulus 1-2, left-right ^5,6^. The linear classifier was trained on pseudo trials built from randomly selected trials. In more detail, for every neuron, we selected 80% of the trials as a training set and the remaining 20% as a testing set to build the pseudo trials. We generated 1000 pseudo trials for training and 1000 pseudo trials for testing per condition. We randomly chose 80% of the training pseudo trials and 20% of the testing pseudo trials to train and test the linear decoder for a total of 100 cross-validations. We showed the final accuracy of the linear decoder as the ratio between the number of correct predictions and the total number of predictions on the testing set averaged across the cross-validations.

To evaluate the statistical significance of the neural signal, we built a null model by randomly shuffling the task condition labels among the pseudo trials. We trained a linear decoder on the shuffled training set for each shuffle and assessed its accuracy on the shuffled testing set. We repeated the shuffle procedure 100 times, obtaining a null model distribution. We defined the chance interval as the interval between 2 standard deviations of the null model distribution around the chance level at 50%. We applied the same procedure above to decode both the neural and artificial data. For the neural data, we decoded the activity in a 200ms time bin stepped by 20ms along time from 200ms before the stimulus onset until 800ms after. We also decoded the neuronal activity from 400ms before the movement onset until 800ms after the response. All decoding analyses were performed using scripts of the scikit-learn SVC package^5^ and custom Python scripts.

##### Model of input signals

In our first model, we modeled the visual and motor input signals to the Purkinje cells differently in early and late learning. During early learning, we modeled the visual input signals for stimuli 1 and 2 as a matrix *V*_1,2_ of (1000 trials, 50 units, 150 time bins) dimensions. At the time bin 0 we set the stimulus onset, and it remains active for all following 150 time bins. Each element of the matrix *V*_1,2_ is defined as follows:

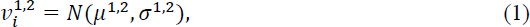

where *i* = 1, . . . ,50 indicates the unit id, *N* is the Gaussian distribution with mean μ^1,2^ equal to - 0.2 and -0.4 for stimulus 1 and stimulus 2, respectively, and a standard deviation σ^1,2^ equal to 0.3 for both stimuli. For each time step and trial, we randomly sample a random value from the previous distributions for each unit.

We modeled the motor signals, i.e., response 1 and 2, as a matrix *M*_1,2_ of (1000 trials, 50 units, 150 time bins) dimension. The movement onset was set at bin 100. Each element *m*_1,2_ of matrices *M*_1,2_ before and after the movement onset is defined as follows:

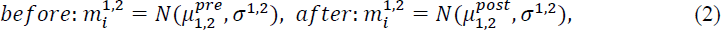

where *i* = 1, . . . ,50 indicates the unit id, *N* is the Gaussian distribution with mean 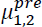 equal to 0 for before the movement onset, and mean 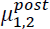 equal to 0.2 and 0.4 for response 1 and response 2 after the movement onset, respectively. The standard deviation σ^1,2^ is set equal to 0.3. In line with the behavioral task, we define two task conditions where visual stimuli 1 (S1) and 2 (S2) are univocally associated with motor responses 1 (R1) and 2 (R2), respectively. When the task condition, for instance, is S1-R1, all the units encoding by construction S2 and R2 are set to not-a-number (nan) during the simulation. We defined a total input matrix *I^input^* of (1000, 200, 150) dimension obtained by concatenating *V*_1,2_ and *M*_1,2_. This input matrix is passed through 100 Rectified Linear Units (ReLU) whose weights *W* are randomly sampled from a Uniform distribution *U*(0,1). The final activity output readout by a linear decoder at time bin *t* is:

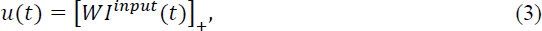

where *I^input^* is the input matrix, and 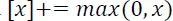 is the ReLU non-linear function mapping the input currents to positive firing rates.

##### Training of an artificial neural network to perform the visuomotor association task

We trained an artificial feedforward neural network to perform the NT and REV tasks. We provided a 5-unit input to the network where a one-hot vector of 3 units encodes the context: the stimulus-only context and task context, i.e., NT or REV tasks. The last one-hot vector of 2 units encodes the stimulus identity. The stimulus identity is always provided to the network along with one of the 3 contexts. The stimulus-only context requires the network to provide the stimulus identity as output regardless of the response (which is random); the task context requires the network still to provide the stimulus identity and the response associated with it according to the NT or REV task.

We introduced the stimulus-only context to reproduce the neural signal dynamics where the visual stimulus is decoded earlier than the response in the late learning phase. In particular, we wanted to model the dynamics of the visual and response neural signals along the trial: in the first three artificial time points, the network receives the stimulus-only context as input to reproduce the onset of the stimulus signal during the experiment. Subsequently, the network receives the task-contextual input and the stimulus identity input, instructing the network on whether to perform the NT or REV task. The stimulus identity is always provided with different contextual inputs at different time points along the trial.

In each time point, the selected input *I* is processed by an expansion layer of 400 linear units through random expansion with initial weights *W^in^* randomly sampled from a uniform distribution *U*(0,1). We refer to these units as artificial Granul Cells (aGCs). The *W^in^* are fixed along the training. The output current α from the aGC unit *k* is defined as the weighted sum of the input as follows:

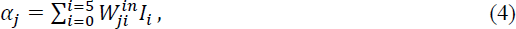

where *j* is the input unit index. The output current is passed to a layer of 100 ReLU units through *W^out^* weights randomly initialized from a uniform distribution *U*(0,1), which will be trained. We refer to these units as artificial Purkinje Cells (aPCs). The output current *β* of an aPC unit *k* is defined as:

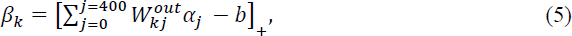

where *j* is the index of an aGC input unit, 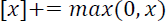 is the non-linear function (ReLu), and *b* is the bias term set equal to 15, and it remains fixed along the training. The output current from aPC layers is read by 2 output units *R*, *S*, which give the stimulus ID and the associated response. The two readouts are defined as follows:

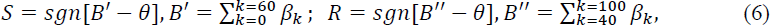

where *θ* = 1 and *B*′ and *B*′′ are the total output currents from the preceding layer’s first 60 and last 60 aPC units. In other words, the two output units *S* and *R* read out the activity from two different pools of aPC units with an overlap of 20 units, i.e., 20%.

The network training is performed over a total of 1000 training trials. In the first 500 trials, the network is trained to receive the stimulus-only and NT-task contexts as input randomly chosen for each trial. In the second half of the training trials, the network is trained to receive the stimulus-only and REV-task contexts. After switching from the first half to the second training phase, we re-initialized the output weights *W^out^* of the last 5% units readout by the stimulus *S* output unit. This is one difference from the actual experiment because we force the network to re-learn the stimulus identity input in order to disambiguate it from the response signal in the following decoding analyses.

The *W^out^* weights are updated in each training trial *t* according to the following rule:

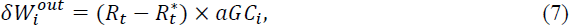

where α*GC_i_* is the output current from the *i*-th aGC unit, *R_t_* is the response the network provides at trial 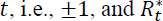 is the correct response. The new output weight is:

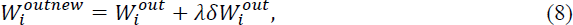

where λ is the learning rate, which is set equal to 0.1 and remains constant throughout the training. The network’s performance in providing the correct stimulus identity and the correct response is tested on a batch of newly generated 100 trials before every training trial. A small amount of noise randomly sampled from a Gaussian distribution of zero mean and variance equal to 10^−3^ is added to the input.

## Acknowledgments

We thank Dr. Rivka Shoulson and Dr. Keely Wharton for excellent veterinary care, Vipul Patel for superb electronic and computer work, John Caban for machining, and Danielle Shank for facilitating everything.

## Author contributions

A.E.I., N.S., and M.E.G. conceived and designed the experiments and collected the data. A.E.I. V.F., S.F., and M.E.G. conceptualized and developed the analyses. A.E.I. developed the single neuron analyses. V.F. and S.F. developed the population analyses and the models. The data were interpreted by A.E.I., V.F., C.D.Z., S.F., and M.E.G., who also wrote the article.

## SUPPLEMENTARY FIGURES

**Supplementary Figure 1.**
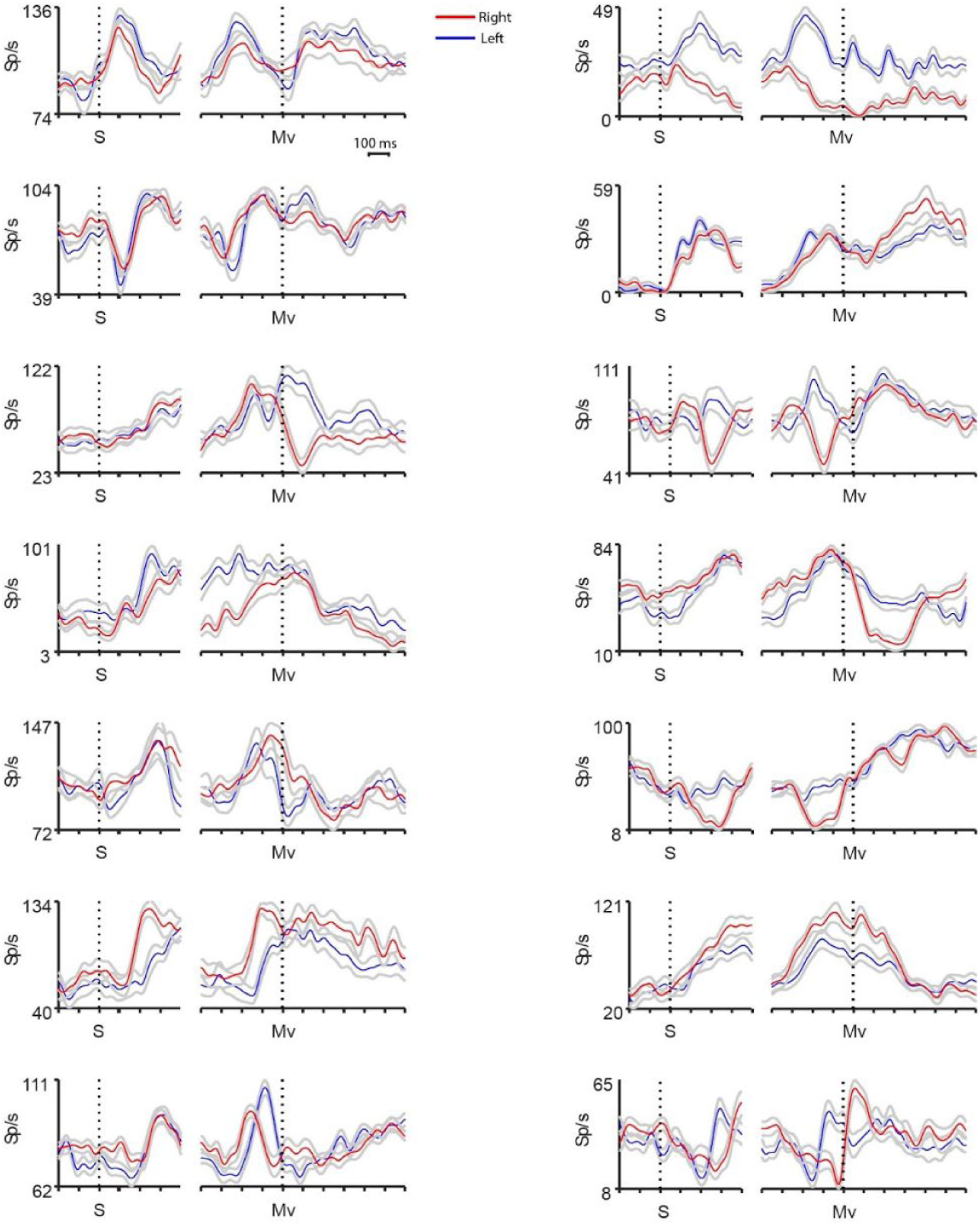
Response of simple spikes in OT. Each panel represents the firing rate (Sp/s) of a single neuron in SR left (blue) and SR right (red) correct trials, aligned wit the onset of the stimulus (S) and with the movement (M).

**Supplementary Figure 2.**
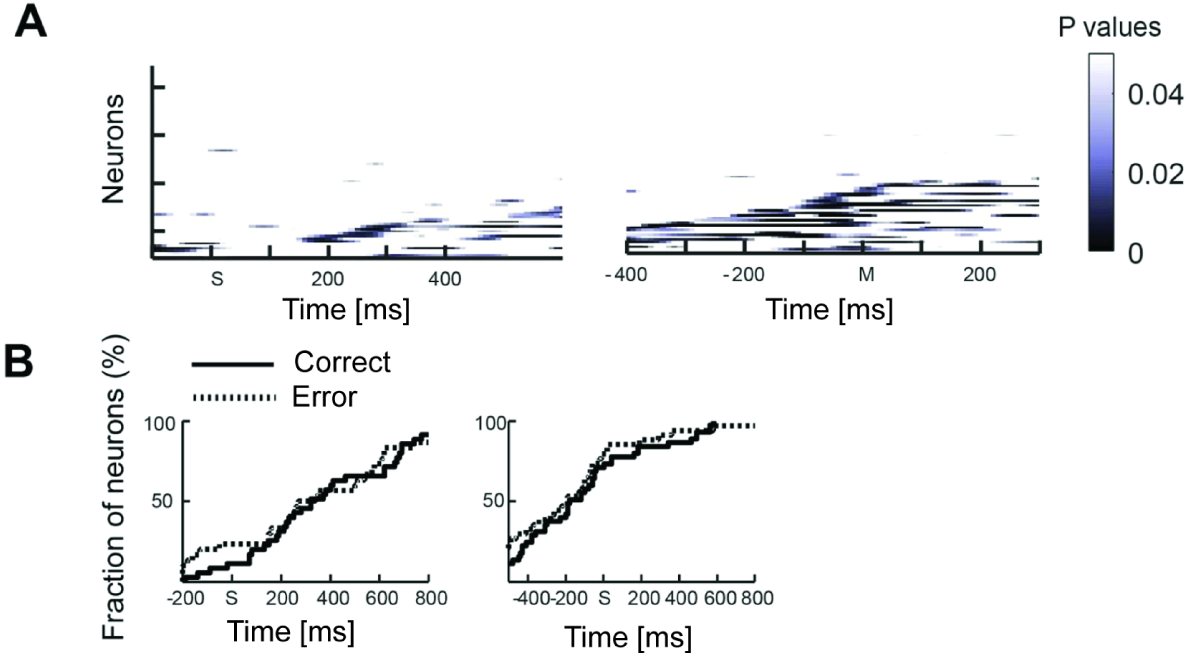
Fraction of response selective neurons in correct and error trials. **A)** Each line represents the p-values between the response to the left and right hand in error trials in early learning for each neuron (y-axis) in 80ms bins, 10ms shifted, from the onset of the stimulus (left) and from the onset of the movement (right) **B)** Empirical cumulative distribution of the times when the population starts to discriminate between the left and the right hand in early learning: in correct trials (solid line) and error trials (dotted line).

**Supplementary Figure 3.**
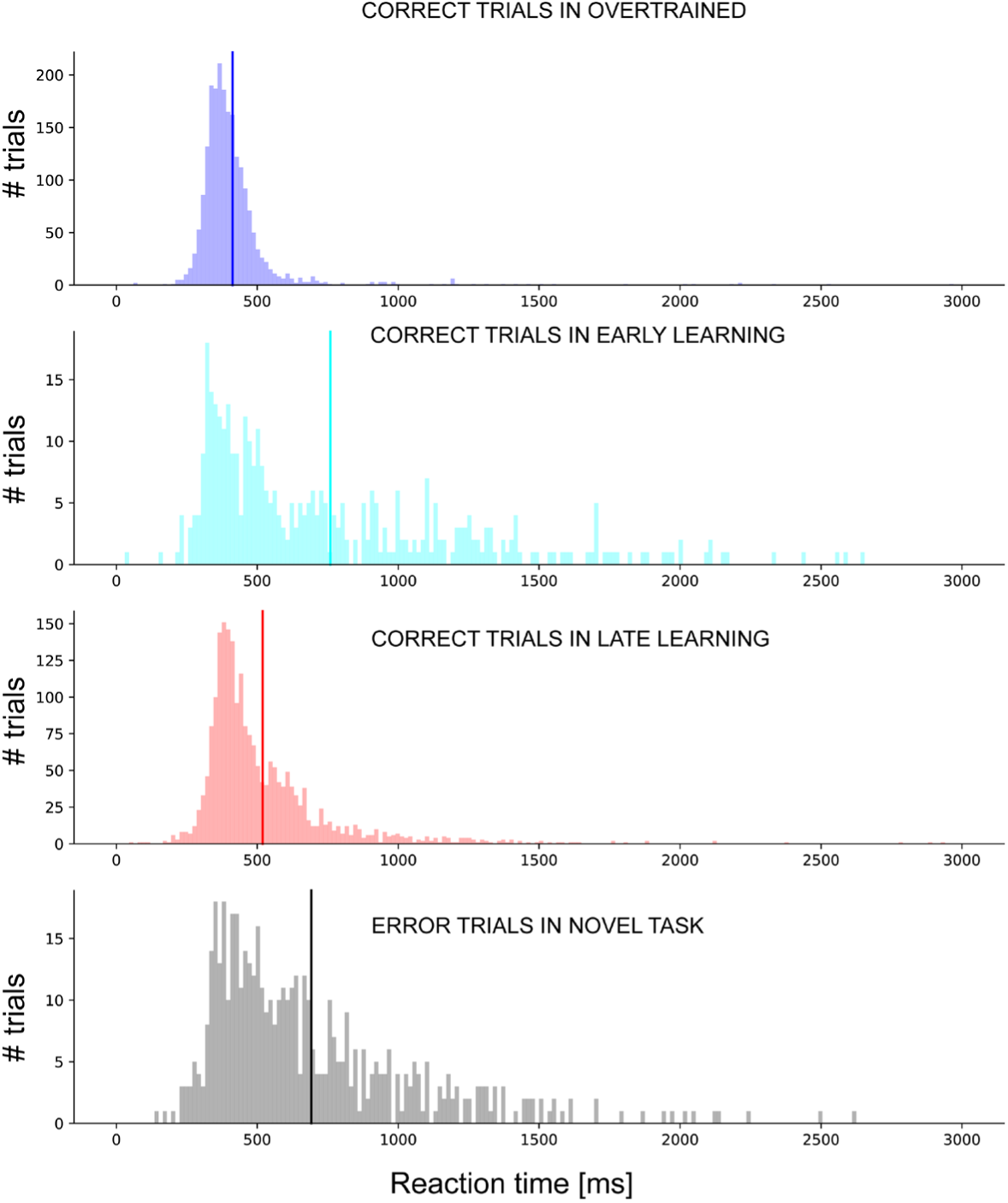
Distribution of reaction times across trials. From top to bottom, distribution of reaction times across trials for correct trials during overtrained (OT) task, early learning (EL) and late learning (LL) in novel tasks (NT), and for error trials during NT. The vertical lines indicate the mean of each distribution. In particular in OT the mean reaction time is (412 ± 170ms) (mean±SD), in EL is (760 ± 484)ms, in LL is (519 ± 243)ms, and in error trials during NT is (691 ± 377)ms. The high value of SD is due to large variability across sessions, mirrored by the skewness of the distributions. SD: standard deviation.

**Supplementary Figure 4.**
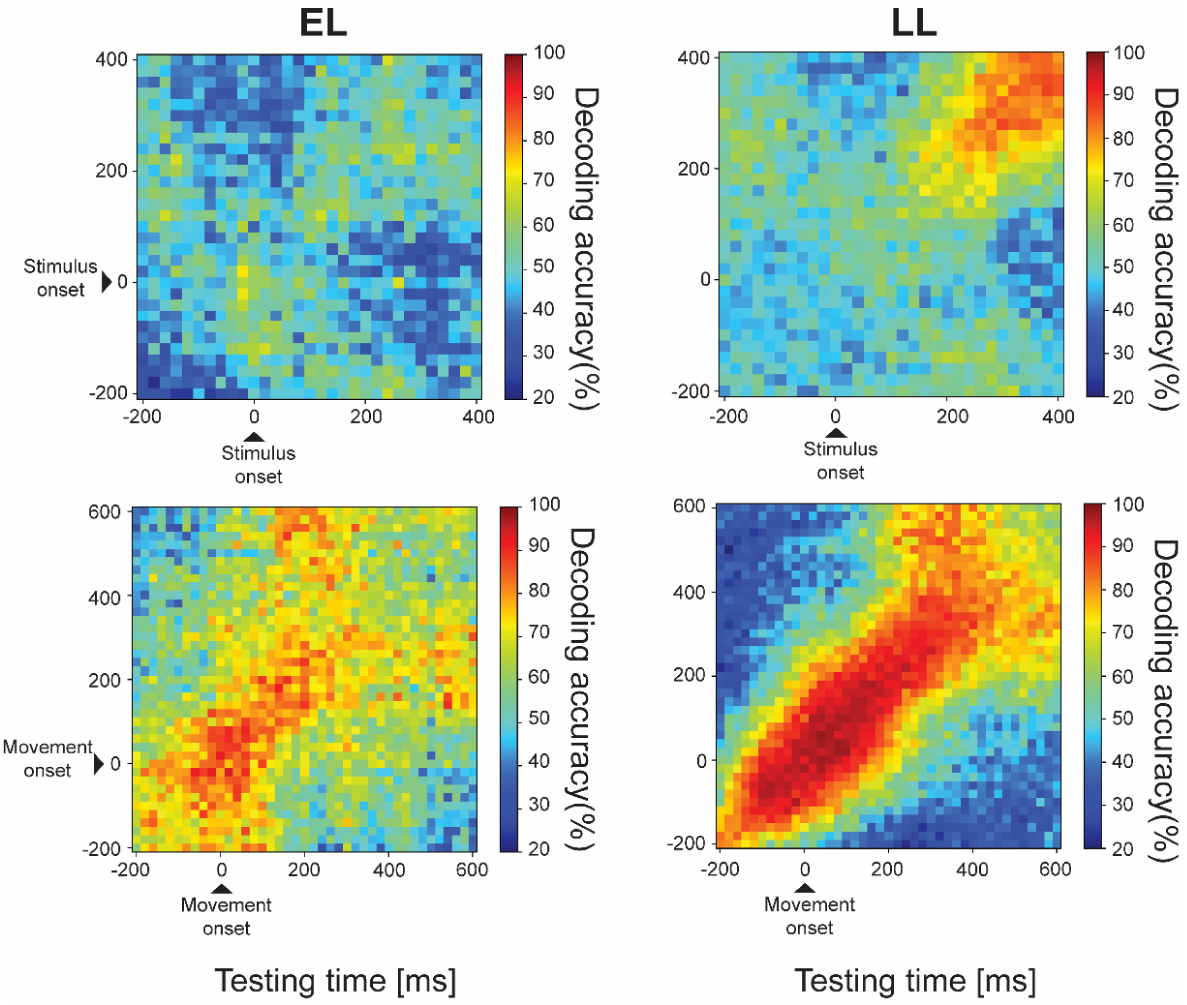
Signal stability analysis in Early learning (EL) and late learning (LL) during Novel Task. Cross-bin decoding where a linear decoder is trained on a time bin and tested on the following or preceding time bins. Left-column: cross-bin decoding analysis during EL around stimulus onset (top) and movement onset (bottom). Right-column: cross-bon decoding analysis in LL around stimulus onset (top) and movement onset (bottom).

**Supplementary Figure 5.**
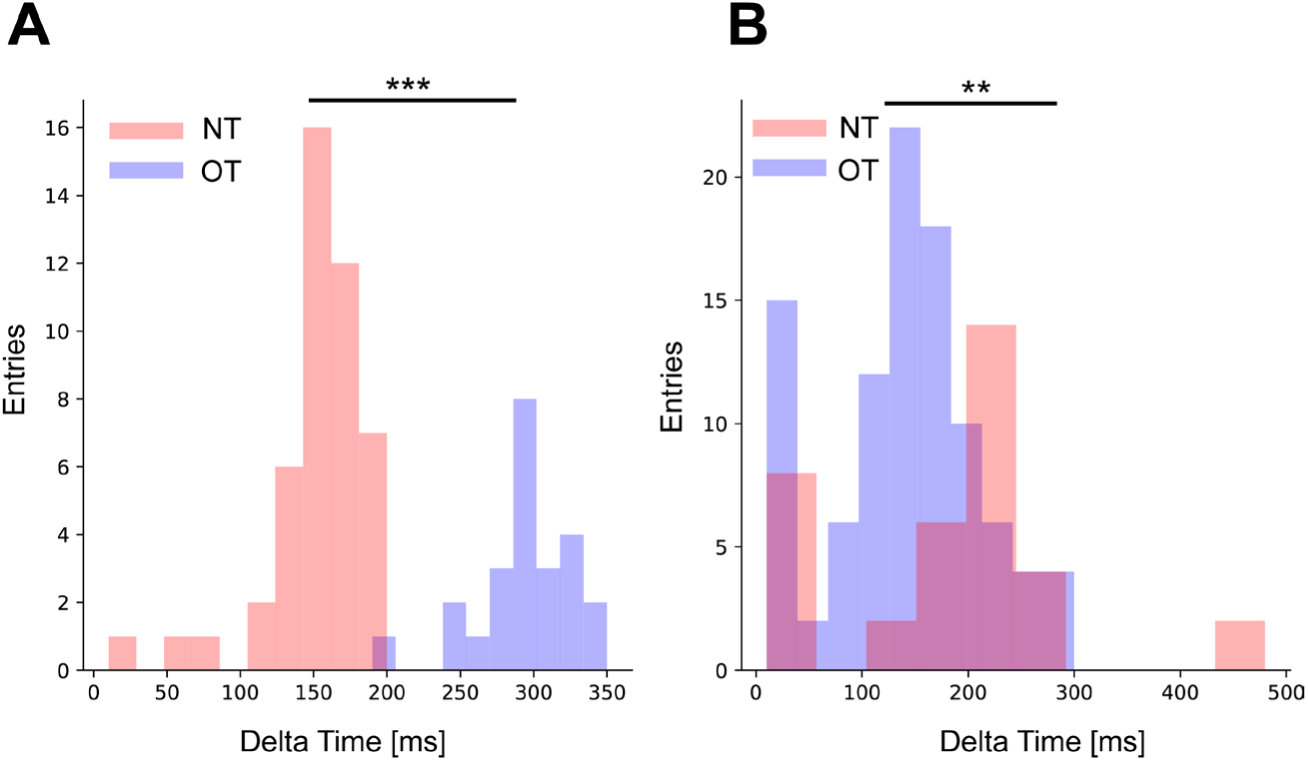
Distribution of time interval where the signal is static. **A)** Distribution of time interval with static representation of the variables after the stimulus onset. The representation is significant more stable in Overtrained Task (OT) than Novel Task (NT) (p-value<0.001; Mann-Whitney U test). **B)** Distribution of time interval with static representation of the variables after the movement onset. The representation is significantly more stable in NT than OT (p-value=0.002; Mann-Whitney U test). *** p-value<0.001, ** p-value <0.01.

**Supplementary Figure 6.**
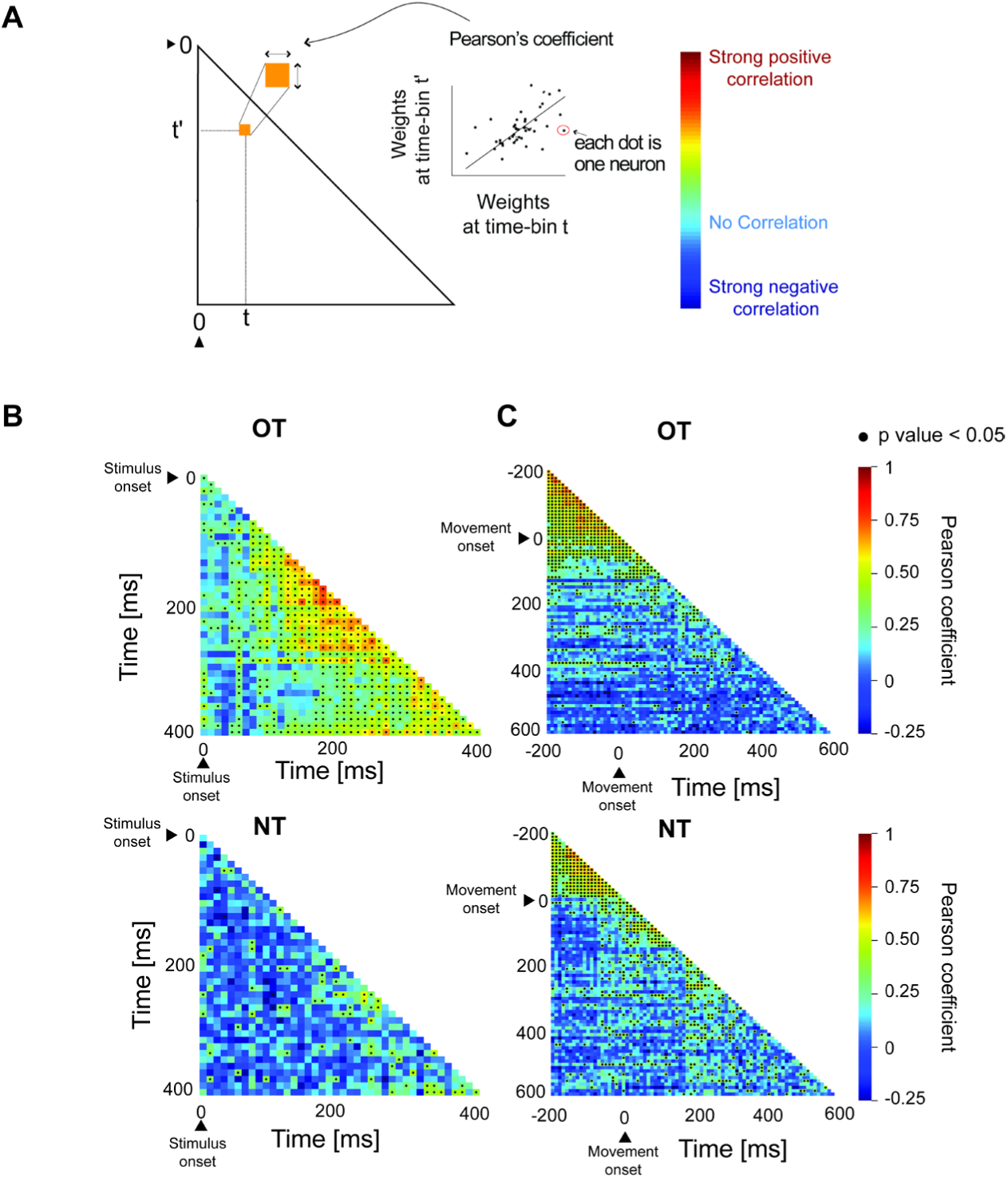
Correlation of the decoder weights along time. **A)** Schematic of the presentation of results in the following panels. **B) Top:** Correlation of the decoder weights between different time bins during OT after the stimulus onset. The higher the Pearson coefficient the more stable is the representation of the variables between different time bins. The black dots indicate where the Pearson coefficient is significant (p-value<0.05). It is evident that the correlation between weights is sustained for several consecutive time bins after the stimulus onset. **Bottom:** Correlation of decoding weights between different time bins in NT after stimulus onset. **C) Top:** Same as in B) around movement onset. **Bottom**: same as in B) around movement onset.

**Supplementary Figure 7.**
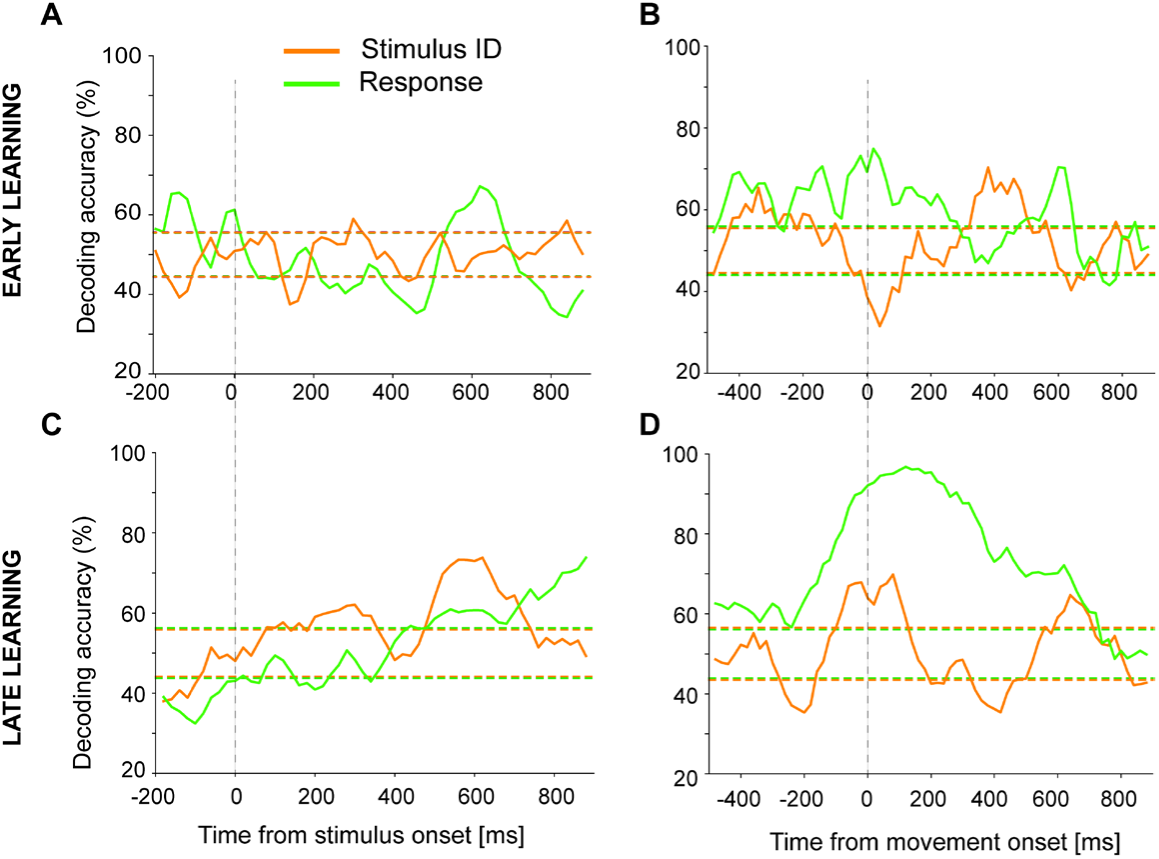
Decoding of the response and stimulus identity signal in correct and error trials in novel task. **A)** Decoding of the response and stimulus identity around the stimulus onset during the early learning phase of the novel task. Correct and error trials have been analyzed to disentangle the two signals. **B)** The same as in A) with time aligned to the movement onset. **C)** Same as in A) during the late learning phase of the novel task. **D)** The same as in C) with time aligned to the movement onset.

## BIBLIOGRAPHY

1 Bastian, A. J. & Lisberger, S. G. Principle of Neuroscience. 6th edn, 908–931 (McGraw Hill, 2021).

2 Sendhilnathan, N., Semework, M., Goldberg, M. E. & Ipata, A. E. Neural Correlates of Reinforcement Learning in Mid-lateral Cerebellum. Neuron 106, 1055, doi:10.1016/j.neuron.2020.05.021 (2020).

3 Sendhilnathan, N., Bostan, A. C., Strick, P. L. & Goldberg, M. E. A cerebro-cerebellar network for learning visuomotor associations. Nature Communication 15, doi:10.1038/s41467-024-46281-0 (2024).

4 Paton, J. J., Belova, M. A., Morrison, S. E. & Salzman, C. D. The primate amygdala represents the positive and negative value of visual stimuli during learning. Nature 439, 865–870, doi:10.1038/nature04490 (2006).

5 Pedregosa, F. et al. Scikit-learn:Machine Learning in Python. Journal of Machine Learning Research 12, 2925–2830 (2011).

6 Bishop, C. M. Patter Recognition and Machine Learning. (Springer, 2006).

7 Asaad, W. F., Rainer, G. & Miller, E. K. Neural activity in the primate prefrontal cortex during associative learning. Neuron 21, 1399–1407 (1998).

8 Pasupathy, A. & Miller, E. K. Different time courses of learning-related activity in the prefrontal cortex and striatum. Nature 433, 873–876, doi:10.1038/nature03287 (2005).

9 Histed, M. H., Pasupathy, A. & Miller, E. K. Learning substrates in the primate prefrontal cortex and striatum: sustained activity related to successful actions. Neuron 63, 244–253, doi:10.1016/j.neuron.2009.06.019 (2009).

10 Gao, Z. et al. A cortico-cerebellar loop for motor planning. Nature 563, 113–116, doi:10.1038/s41586-018-0633-x (2018).

11 Chabrol, F. P., Blot, A. & Mrsic-Flogel, T. D. Cerebellar Contribution to Preparatory Activity in Motor Neocortex. Neuron 103, 506–519 e504, doi:10.1016/j.neuron.2019.05.022 (2019).

12 Yang, Y. & Lisberger, S. G. Role of plasticity at different sites across the time course of cerebellar motor learning. J Neurosci 34, 7077–7090, doi:10.1523/JNEUROSCI.0017-14.2014 (2014).

13 ten Brinke, M. M. et al. Evolving Models of Pavlovian Conditioning: Cerebellar Cortical Dynamics in Awake Behaving Mice. Cell Rep 13, 1977–1988, doi:10.1016/j.celrep.2015.10.057 (2015).

14 Ohmae, S. & Medina, J. F. Climbing fibers encode a temporal-difference prediction error during cerebellar learning in mice. Nat Neurosci 18, 1798–1803, doi:10.1038/nn.4167 (2015).

15 Voges, K., Wu, B., Post, L., Schonewille, M. & De Zeeuw, C. I. Mechanisms underlying vestibulo-cerebellar motor learning in mice depend on movement direction. J Physiol 595, 5301–5326, doi:10.1113/JP274346 (2017).

16 Sendhilnathan, N., Ipata, A. & Goldberg, M. E. Mid-lateral cerebellar complex spikes encode multiple independent reward-related signals during reinforcement learning. Nat Commun 12, 6475, doi:10.1038/s41467-021-26338-0 (2021).

17 Giovannucci, A. et al. Cerebellar granule cells acquire a widespread predictive feedback signal during motor learning. Nat Neurosci 20, 727–734, doi:10.1038/nn.4531 (2017).

18 Broersen, R. et al. Synaptic mechanisms for associative learning in the cerebellar nuclei. Nat Commun 14, 7459, doi:10.1038/s41467-023-43227-w (2023).

19 Mello, G. B., Soares, S. & Paton, J. J. A scalable population code for time in the striatum. Curr Biol 25, 1113–1122, doi:10.1016/j.cub.2015.02.036 (2015).

20 de Zeeuw, C. I. & Berrebi, A. S. Individual Purkinje cell axons terminate on both inhibitory and excitatory neurons in the cerebellar and vestibular nuclei. Ann N Y Acad Sci 781, 607–610, doi:10.1111/j.1749-6632.1996.tb15736.x (1996).

21 De Zeeuw, C. I. & Berrebi, A. S. Postsynaptic targets of Purkinje cell terminals in the cerebellar and vestibular nuclei of the rat. Eur J Neurosci 7, 2322–2333, doi:10.1111/j.1460-9568.1995.tb00653.x (1995).

22 Petajan, J. H. AAEM minimonograph #3: motor unit recruitment. Muscle Nerve 14, 489–502, doi:10.1002/mus.880140602 (1991).

23 Xiao, J. et al. Systematic regional variations in Purkinje cell spiking patterns. PLoS One 9, e105633, doi:10.1371/journal.pone.0105633 (2014).

24 Zhou, H. et al. Cerebellar modules operate at different frequencies. Elife 3, e02536, doi:10.7554/eLife.02536 (2014).

25 Cerminara, N. L., Lang, E. J., Sillitoe, R. V. & Apps, R. Redefining the cerebellar cortex as an assembly of non-uniform Purkinje cell microcircuits. Nat Rev Neurosci 16, 79–93, doi:10.1038/nrn3886 (2015).

26 Wu, B. et al. TRPC3 is a major contributor to functional heterogeneity of cerebellar Purkinje cells. Elife 8, doi:10.7554/eLife.45590 (2019).

27 De Zeeuw, C. I., Koppen, J., Bregman, G. G., Runge, M. & Narain, D. Heterogeneous encoding of temporal stimuli in the cerebellar cortex. Nat Commun 14, 7581, doi:10.1038/s41467-023-43139-9 (2023).

28 Thier, P., Dicke, P. W., Haas, R. & Barash, S. Encoding of movement time by populations of cerebellar Purkinje cells. Nature 405, 72–76, doi:10.1038/35011062 (2000).

29 Herzfeld, D. J., Kojima, Y., Soetedjo, R. & Shadmehr, R. Encoding of action by the Purkinje cells of the cerebellum. Nature 526, 439–442, doi:10.1038/nature15693 (2015).

30 Hong, S. et al. Multiplexed coding by cerebellar Purkinje neurons. Elife 5, doi:10.7554/eLife.13810 (2016).

31 Li, J. X., Medina, J. F., Frank, L. M. & Lisberger, S. G. Acquisition of neural learning in cerebellum and cerebral cortex for smooth pursuit eye movements. J Neurosci 31, 12716–12726, doi:10.1523/JNEUROSCI.2515-11.2011 (2011).

32 Dash, S., Dicke, P. W. & Thier, P. A vermal Purkinje cell simple spike population response encodes the changes in eye movement kinematics due to smooth pursuit adaptation. Front Syst Neurosci 7, 3, doi:10.3389/fnsys.2013.00003 (2013).

33 Zobeiri, O. A. & Cullen, K. E. Cerebellar Purkinje cells in male macaques combine sensory and motor information to predict the sensory consequences of active self-motion. Nat Commun 15, 4003, doi:10.1038/s41467-024-48376-0 (2024).

34 De Zeeuw, C. I. et al. Spatiotemporal firing patterns in the cerebellum. Nat Rev Neurosci 12, 327–344, doi:10.1038/nrn3011 (2011).

35 Huang, C. C. et al. Convergence of pontine and proprioceptive streams onto multimodal cerebellar granule cells. Elife 2, e00400, doi:10.7554/eLife.00400 (2013).

36 Ishikawa, T., Shimuta, M. & Häusser, M. Multimodal sensory integration in single cerebellar granule cells in vivo. Elife 4, doi:10.7554/eLife.12916 (2015).

37 Galliano, E. et al. Silencing the majority of cerebellar granule cells uncovers their essential role in motor learning and consolidation. Cell Rep 3, 1239–1251, doi:10.1016/j.celrep.2013.03.023 (2013).

38 Wagner, M. J., Kim, T. H., Savall, J., Schnitzer, M. J. & Luo, L. Cerebellar granule cells encode the expectation of reward. Nature 544, 96–100, doi:10.1038/nature21726 (2017).

39 Guo, C. et al. Purkinje Cells Directly Inhibit Granule Cells in Specialized Regions of the Cerebellar Cortex. Neuron 91, 1330–1341, doi:10.1016/j.neuron.2016.08.011 (2016).

40 Witter, L., Rudolph, S., Pressler, R. T., Lahlaf, S. I. & Regehr, W. G. Purkinje Cell Collaterals Enable Output Signals from the Cerebellar Cortex to Feed Back to Purkinje Cells and Interneurons. Neuron 91, 312–319 (2016).

41 Boyden, E. S., Katoh, A. & Raymond, J. L. Cerebellum-dependent learning: the role of multiple plasticity mechanisms. Annu Rev Neurosci 27, 581–609, doi:10.1146/annurev.neuro.27.070203.144238 (2004).

42 Gao, Z., van Beugen, B. J. & De Zeeuw, C. I. Distributed synergistic plasticity and cerebellar learning. Nat Rev Neurosci 13, 619–635, doi:10.1038/nrn3312 (2012).

43 Sendhilnathan, N., Goldberg, M. E. & Ipata, A. E. Mixed Selectivity in the Cerebellar Purkinje-Cell Response during Visuomotor Association Learning. J Neurosci 42, 3847–3855, doi:10.1523/JNEUROSCI.1771-21.2022 (2022).

44 Rigotti, M. et al. The importance of mixed selectivity in complex cognitive tasks. Nature 497, 585–590, doi:10.1038/nature12160 (2013).

